# Development of the zebrafish anterior lateral line system is influenced by underlying cranial neural crest

**DOI:** 10.1101/2025.02.11.637483

**Authors:** Vishruth Venkataraman, Noel H. McGrory, Theresa J. Christiansen, Joaquin Navajas Acedo, Michael I. Coates, Victoria E. Prince

## Abstract

The mechanosensory lateral line system of aquatic vertebrates comprises a superficial network of distributed sensory organs, the neuromasts, which are arranged over the head and trunk and innervated by lateral line nerves to allow detection of changes in water flow and pressure. While the well-studied zebrafish posterior lateral line has emerged as a powerful model to study collective cell migration, far less is known about development of the anterior lateral line, which produces the supraorbital and infraorbital lines around the eye, as well as mandibular and opercular lines over the jaw and cheek. Here we show that normal development of the zebrafish anterior lateral line system from cranial placodes is dependent on another vertebrate-specific cell type, the cranial neural crest. We find that cranial neural crest and anterior lateral lines develop in close proximity, with absence of neural crest cells leading to major disruptions in the overlying anterior lateral line system. Specifically, in the absence of neural crest neither supraorbital nor infraorbital lateral lines fully extend, such that the most anterior cranial regions remain devoid of neuromasts, while supernumerary ectopic neuromasts form in the posterior supraorbital region. Both neural crest and cranial placodes contribute neurons to the lateral line ganglia that innervate the neuromasts and in the absence of neural crest these ganglia, as well as the lateral line afferent nerves, are disrupted. Finally, we establish that as ontogeny proceeds, the most anterior supraorbital neuromasts come to lie within neural crest-derived frontal and nasal bones in the developing cranium. These are the same anterior supraorbital neuromasts that are absent or mislocated in specimens lacking neural crest cells. Together, our results establish that cranial neural crest and cranial placode derivatives function in concert over the course of ontogeny to build the complex cranial lateral line system.

**Highlights:**

- The anterior lateral line and cranial neural crest develop in close proximity
- Absence of neural crest disrupts anterior lateral line development
- Absence of neural crest disrupts lateral line ganglion morphology and innervation
- Early interactions of neural crest and placodes prefigure later anatomical interactions

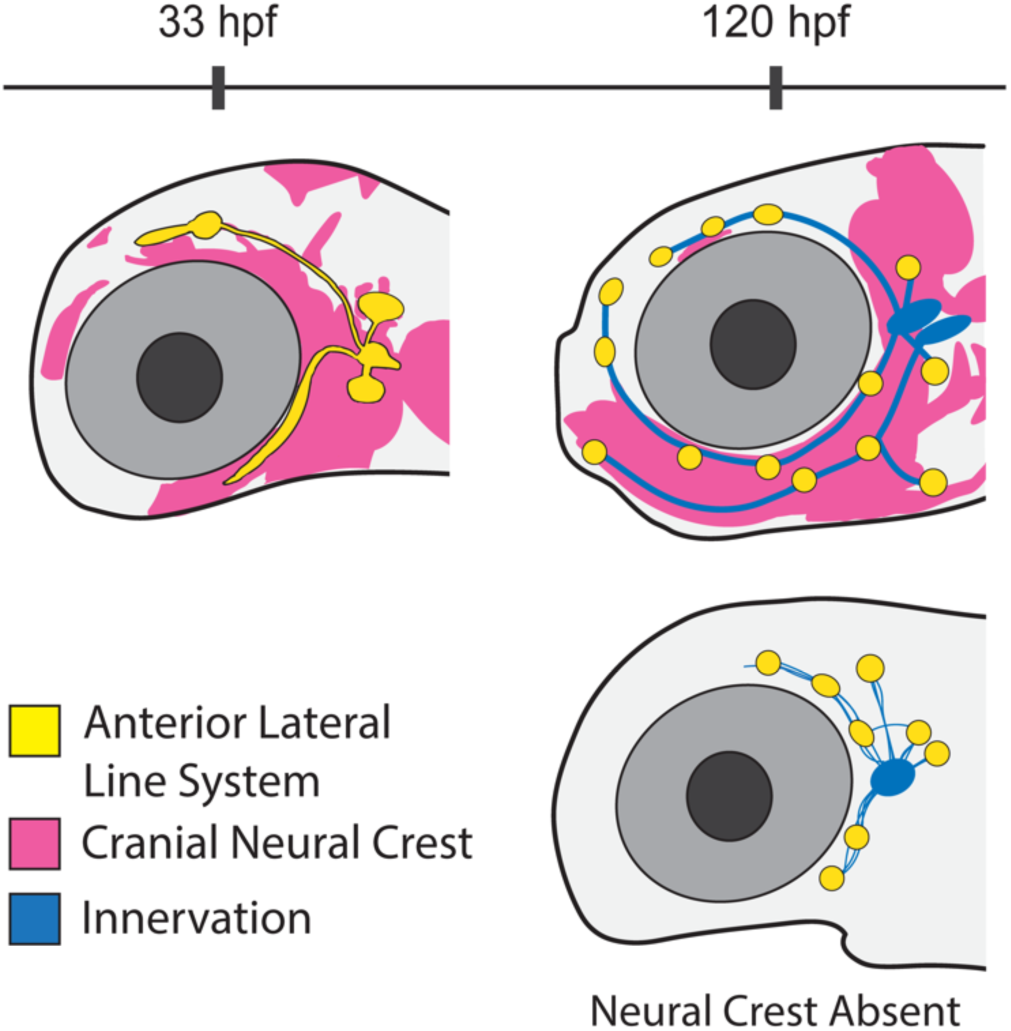

## 1. Introduction

Aquatic vertebrates possess a mechanosensory lateral line system that gives organisms the ability to detect water movements in their environment. The lateral line system serves to orient an individual relative to flow direction, facilitate schooling behavior, enhance prey location, and contribute to predator evasion (Mogdans, 2019). Recent functional studies in goldfish have shown that this mechanosensory system is part of a broader multimodal ‘bioacoustic’ network, working together with the ear to detect auditory signals in the water (Higgs & Radford, 2013). The system includes the neuromasts—each comprising a core of sensory hair cells surrounded by support cells—and their afferent neurons, both of which are derived embryologically from discrete patches of thickened neurogenic epithelium termed lateral line placodes. The lateral line placodes form bilaterally on the sides of the developing head and each gives rise to a primordium and a sensory ganglion, which differentiate into the sense organs and nerves of the lateral line system, respectively.

The cellular and molecular bases of zebrafish posterior lateral line development have been studied in detail (reviewed by Chitnis et al., 2012; Olson & Nechiporuk, 2018; Piotrowski & Baker, 2014). Briefly, the posterior lateral line initially develops from a post-otic primordium which migrates caudally along the trunk as a dynamic collective, depositing neuromasts in its wake at defined locations. At the same time, the primordium tows along its afferent nerve, which extends from the posterior lateral line ganglion (Haas & Gilmour, 2006). Key roles in this complex process have been established for chemokine signaling, which helps mediate directed migration, and for Fgf and Wnt signaling, which coordinate primordium proliferation and neuromast deposition (reviewed by Piotrowski & Baker, 2014). Posterior lateral line development continues as a second primordium follows the first, depositing additional neuromasts (Navajas Acedo et al., 2019), with intercalary neuromasts emerging still later, through proliferation of interneuromast cells left behind by the migrating primordia (reviewed by Ghysen & Dambly-Chaudière, 2007).

We have a more limited understanding of zebrafish anterior lateral line development, derived primarily from two previous studies. A description of the anterior lateral line ganglia and nerves was published over two decades ago (Raible & Kruse, 2000) and, more recently, transgenic zebrafish lines were used to generate a detailed description of the deposition of anterior lateral line neuromasts, and their innervation, over the first 10 days of development (Iwasaki et al., 2020). We also know that the initial establishment of the placodes that form posterior versus anterior lateral line primordia is controlled by different signaling pathways: retinoic acid is necessary for formation of the posterior lateral line placode, while Fgf is necessary for formation of the anterior lateral line placode (Nikaido et al., 2017). While disruptions in chemokine signaling block migration of the posterior lateral line, they have not been reported to disrupt the anterior lateral line (David et al., 2002; Ghysen & Dambly-Chaudière, 2007).

The anterior lateral lines that extend above the eye (the supraorbital) and below the eye (the infraorbital) develop from an anterodorsal placode, which arises just anterior to the otic vesicle by 24 hours post fertilization (hpf). The lines that traverse the cheek and jaw develop from an anteroventral placode, which arises a little more ventrally around 36 hpf (Iwasaki et al., 2020; Raible & Kruse, 2000). Iwasaki and colleagues (2020) classified anterior lateral line neuromasts into four categories: homegrown, which develop near their point of origin; migratory, which arise from a migratory primordium; budding, which form through outgrowth from already deposited migratory neuromasts; and intercalary, which arise from interneuromast cells, similar to the production of intercalary neuromasts in the posterior lateral line.

Cranial placodes, together with neural crest cells, are found exclusively in the vertebrates and these tissues have together allowed the evolution of complex vertebrate crania with their array of sensory structures (Gans & Northcutt, 1983). These two classes of cells develop from early primordia that lie in close apposition and are likely intermixed at the earliest stages (reviewed by Koontz et al., 2023; Rocha, Beiriger, et al., 2020; Rocha, Singh, et al., 2020; Steventon et al., 2014). As development continues, placode-derived and neural crest cells remain close together, allowing scope for interactions between these spatially associated cells in the assembly of cranial sensory structures (Steventon et al., 2014). Cranial neural crest cells are therefore a good candidate to influence the development of the anterior lateral line.

Functional interplay between neural crest and cranial placodes has been described in multiple species and situations, including in zebrafish posterior lateral line development, where neural crest-derived glial cells function to suppress premature formation of trunk intercalary neuromasts (Grant et al., 2005; López-Schier & Hudspeth, 2005; Lush & Piotrowski, 2014). In chick, as well as zebrafish, both neural crest and placode cells co-contribute to cranial ganglia (Covell Jr. & Noden, 1989; Hamburger, 1961; Kague et al., 2012). Chick neural crest cells also establish corridors for placode-derived neuroblast cells to migrate through, helping to establish stereotypical organization of their cranial sensory ganglia (Freter et al., 2013). Zebrafish neural crest cells similarly promote assembly of epibranchial ganglia (Culbertson et al., 2011).

Evidence of neural crest-placode interactions has also been found in a lamprey, *Petromyzon*, where CRISPR-mediated disruption of neural crest development results in changes to cranial ganglia morphology and positioning, but does not affect the total number of ganglia (Yuan et al., 2020). In *Xenopus*, the positioning of both the neural crest cells and the placodes is partially determined by a ‘chase and run’ dynamic. The neural crest cells ‘chase’ placode cells due to attraction by placode-secreted chemokines, whereas placode cells ‘run’ from neural crest cells in response to cell contact (Theveneau et al., 2013). Importantly, the zebrafish neural crest and anterior lateral line systems remain closely coupled throughout the entire course of ontogeny. The cranial neural crest gives rise to a subset of the zebrafish dermal bones that encase the adult cranium (Kague et al., 2012), and as bones form during ontogeny, the cranial lateral lines become enclosed by bony canals (Webb & Shirey, 2003).

The relationship between the lateral line and surrounding dermal bone was a topic of repeated interest over the last century, with multiple authors suggesting that lateral line neuromasts might influence ossification of surrounding dermal bones (Moy-Thomas, 1947; Pehrson, 1922; Stensiö, 1947). In response to that idea, but also to findings that called it into question, Parrington (1949) put forward an alternative model rooted in early development. Specifically, Parrington hypothesized that “the precursors of dermal ossifications influence the courses of the lateral lines”. While Parrington did not mention neural crest, we now know that the neural crest cells are the source of many cranial dermal bones (Kague et al., 2012). Parrington’s hypothesis, together with the many examples of functional interactions between neural crest and cranial placodes (above), led us to test whether zebrafish neural crest cells might interact with the placode-derived anterior lateral line system to influence how the lateral lines navigate over the developing head.

Here, we have investigated zebrafish anterior lateral line development using confocal and Single Plane Illumination Microscopy (SPIM) of transgenic specimens, as well as immunolabeling, to build on previous descriptions of neuromast deposition and innervation (Iwaskaki et al., 2020, Raible & Kruse, 2000). Importantly, we have established that both the supraorbital and infraorbital anterior lateral lines develop directly adjacent to cranial neural crest cells, with neural crest cells also surrounding and contributing to anterior lateral line ganglia. To test the hypothesis that cranial neural crest influences anterior lateral line development we ablated neural crest cells using transgene mediated cytotoxicity. We found that absence of neural crest cells leads to absence of neuromasts in the most anterior (pre-orbital) part of the cranium. Supraorbital line extension stalls above the eye, while ectopic neuromasts often form in the posterior supraorbital region. The infraorbital line is severely reduced, and the mandibular and opercular lines are missing. We also explored the innervation of the zebrafish anterior lateral line system, demonstrating that both dorsal and ventral anterior lateral line ganglia have neural crest and placode cell contributions, and establishing that lateral line gangliogenesis and lateral line innervation are disrupted in the absence of neural crest cells. Finally, we exploited Cre-based lineage tracing of neural crest cells to reveal that the most anterior supraorbital neuromasts become encased in neural crest-derived bone. These same most anterior supraorbital neuromasts are the ones that fail to migrate to their normal positions in the absence of neural crest. Together, our findings establish that developmental interactions between neural crest and placode-derived anterior lateral line cells prefigure anatomical interactions.

## 2. Results

### 2.1 The anterior lateral line system develops in close proximity to cranial neural crest

Throughout this study we have made use of *Tg(cldnB:GFP)* to visualize the dynamic process of anterior lateral line development. This tight junction marker labels the membranes of sensory placode cells including those of the lateral line system, as well as epithelial cells (Haas & Gilmour, 2006). Figure 1A-E shows a series of still images from a confocal time-lapse analysis of a triple transgenic *Tg(cldnB:GFP;dRA:GFP;cldnB:H2AmCherry*) specimen. The *Tg(dRA:GFP*) line (provided by Parker & Krumlauf) is a fortuitous insertion, which labels the lateral line (yellow) and augments the *cldnB:GFP* signal (also yellow); nuclear localized *cldnB:H2AmCherry* (magenta) reveals the nuclei of lateral line and epithelial cells. Our analysis focused on the cranial region, specifically the region between the developing eye and the otic vesicle, from 25-42 hpf, with confocal maximum projection images shown in lateral view (Fig. 1A-E; Supplemental Movies 1, 2). As early as 25 hpf the primordia that will produce the supraorbital (SO) and infraorbital (IO) anterior lateral lines can be observed beginning to migrate anterodorsally and anteroventrally, respectively (Fig. 1A, A’, SOp and IOp). Shortly after this stage, a little before 29 hpf, the first two otic neuromasts are deposited (Fig. 1B, B’, O1 and O2). These ‘homegrown’ neuromasts (Iwasaki et al., 2020) remain close to their points of origin, gradually shifting apart as development continues. At this same 28.7 hpf stage, the IO primordium consists of an elongated ridge of cells extending under the eye (Fig. 1B, B’, IOp). The SO primordium deposits the SO2 neuromast at about 33 hpf (Fig. 1C, C’), and shortly after begins to bud, such that the SO line extends further anteriorly over and around the eye as development proceeds (Fig. 1D-E; and see also Fig. 1G-I’, below). The IO primordium is maintained as a ridge, but gradually starts to condense to form neuromasts (e.g. Fig. 1E, IO4; and see also Fig. 1F-I’, below).

**Figure 1:**
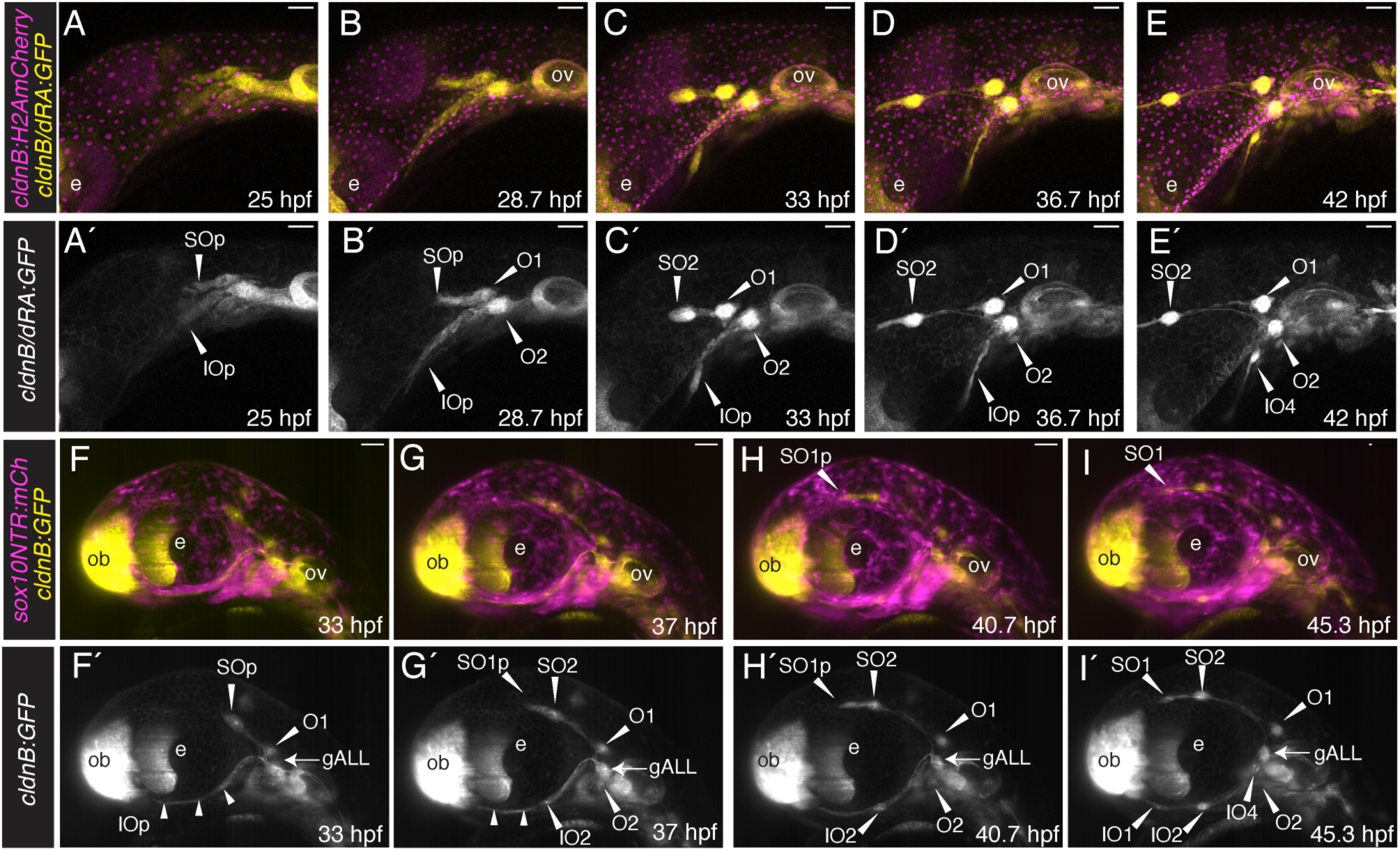
Time-lapse microscopy analysis reveals dynamics of anterior lateral line development and relationship to cranial neural crest cells. (**A-E**) Confocal microscopy time-lapse images of the cranial region of a *Tg(cldnB:GFP;dRA:GFP;cldnB:H2A-mCherry)* triple transgenic specimen, in lateral view with anterior to the left. Maximum projections are shown at 25 hpf (A), 28.7 hpf (B), 33 hpf (C), 36.7 hpf (D), and 42 hpf (E). **A’-E’** are single channel images of the lateral line (*cldnB:GFP* and *dRA:GFP*) alone. Neuromasts are indicated with white arrowheads and labeled (SO: supraorbital neuromast; IO: infraorbital neuromast; O: otic neuromast). The supraorbital and infraorbital primordia are indicated (SOp, IOp). (**F-I**) Single Plane Illumination Microscopy (SPIM) time-lapse images of the cranial region of a *Tg(sox10:NTRmCherry;cldnB:GFP)* double transgenic specimen, in lateral view with anterior to the left, allowing comparison of the development of the lateral line system (*cldnB:GFP*; yellow) and neural crest (*sox10:NTRmCh*; magenta) over developmental time. Maximum projections (imaged using the 20x objective) are shown at 33 hpf (F), 37 hpf (G), 40.7 hpf (H), and 46.3 hpf (I). **F’-I’** are single channel images of the lateral line (*cldnB:GFP*) alone. Neuromasts are indicated with white arrowheads and labeled. Small white arrowheads in F’ and G’ indicate the elongating ridge of the infraorbital line, which can be seen condensing to form the IO2 neuromast in G’-I’. Note budding of the SO2 neuromast in G’ and H’, extending a primordium (SO1p) that then forms the SO1 neuromast, I’. Abbreviations are as follows:-e: lens of the eye; ov: otic vesicle; ob: olfactory bulb. Scale bars are 50 µm.

To investigate a potential relationship between the developing anterior lateral line system and the cranial neural crest, we made use of *Tg(sox10:NTRmCherry)*, which—in common with other transgenes driven by *sox10* gene regulatory sequences—labels developing neural crest cells (Rosenberg et al., 2014). Figure 1F-I shows a series of still maximum projection images from a Single Plane Illumination Microscopy (SPIM) time-lapse analysis of a *Tg(cldnB:GFP;sox10:NTRmCherry)* double transgeni*c s*pecimen. Our SPIM analysis allowed the entire cranial region to be imaged from 33 to 52 hpf, again presented in lateral view (Fig. 1F-1; and Supplemental Videos 3, 4). At 33 hpf the supraorbital primordium of the anterior lateral line is already migrating anterodorsally above the eye (Fig. 1F, F’, SOp). By 37 hpf the SO2 neuromast has started to condense from the supraorbital primordium (Fig. 1G, G’, SO2), and has begun to form SO1 (here labeled as SO1p, for primordium). The budding process continues (Fig. 1H, H’), depositing the more anteriorly located SO1 neuromast close to the olfactory bulb (ob) by 45 hpf (Fig. 1I, I’). The infraorbital primordium, in contrast to the supraorbital, initially forms as an elongated ridge of cells (Fig. 1F, F’, small arrowheads). The IO neuromasts gradually condense from this ridge, with IO2 beginning to appear by 37 hpf (Fig. 1G, G’), and the more anteriorly located IO1 by 45 hpf (Fig. 1I, I’). Over this same developmental period the homegrown otic neuromasts O1 and O2, remain dorsal and ventral to the anterior lateral line ganglion (gALL; which is also cldnB:GFP-positive), gradually shifting apart as development proceeds. Throughout the 33-52 hpf developmental time-period, the various lateral line components are in close proximity to neural crest cells that have already migrated out into the cranial region (Fig. 1F-I; Supplemental Video 4). Above the eye, the region into which SO2 buds to produce SO1 houses a layer of abundant neural crest cells (Fig. 1H; I; note locations of SO1p and SO1). Below the eye, where the IO line is developing, a dense domain of neural crest cells is present throughout the region (Fig. 1F-I’).

In summary, confirming previous descriptions (Iwasaki et al., 2020), our confocal and SPIM time-lapse analyses demonstrate that the supraorbital lateral line forms via migration and budding of neuromasts. By contrast, we find that the infraorbital line forms from an initial elongated ridge of cells, with neuromasts then condensing from the ridge. Importantly, our analysis also reveals that anterior lateral line components are already in close proximity to cranial neural crest cells at the 33 hpf stage, a relationship that continues as the IO and SO lines continue to develop, including during the stages when the SO2 neuromast buds to generate SO1.

To investigate the proximity of the anterior lateral line and cranial neural crest cells in more detail we returned to confocal microscopy. We once again used the *Tg(cldnB:GFP)* line to label the lateral line, but for this analysis used *Tg(sox10:mRFP)*, which labels the membranes of neural crest cells (Kucenas et al., 2008), allowing more effective evaluation of cell-cell interactions between neural crest and placode-derived lateral line cells (Figure 2). Confocal imaging of 37 hpf specimens (Fig. 2A, A’) again demonstrates that cells of the placode-derived lateral line system (*cldnB:GFP*, yellow) and neural crest (*sox10:mRFP*, magenta) are closely apposed. In Fig. 2A’ the supraorbital and infraorbital regions are boxed, indicating the regions imaged at higher magnification in panels B and E, respectively. Fig. 2B shows the developing supraorbital region, with boxed regions on this panel indicating the locations of images focused on the SO1 neuromast primordium (C) and SO2 neuromast (D), displayed in subsequent panels.

**Figure 2:**
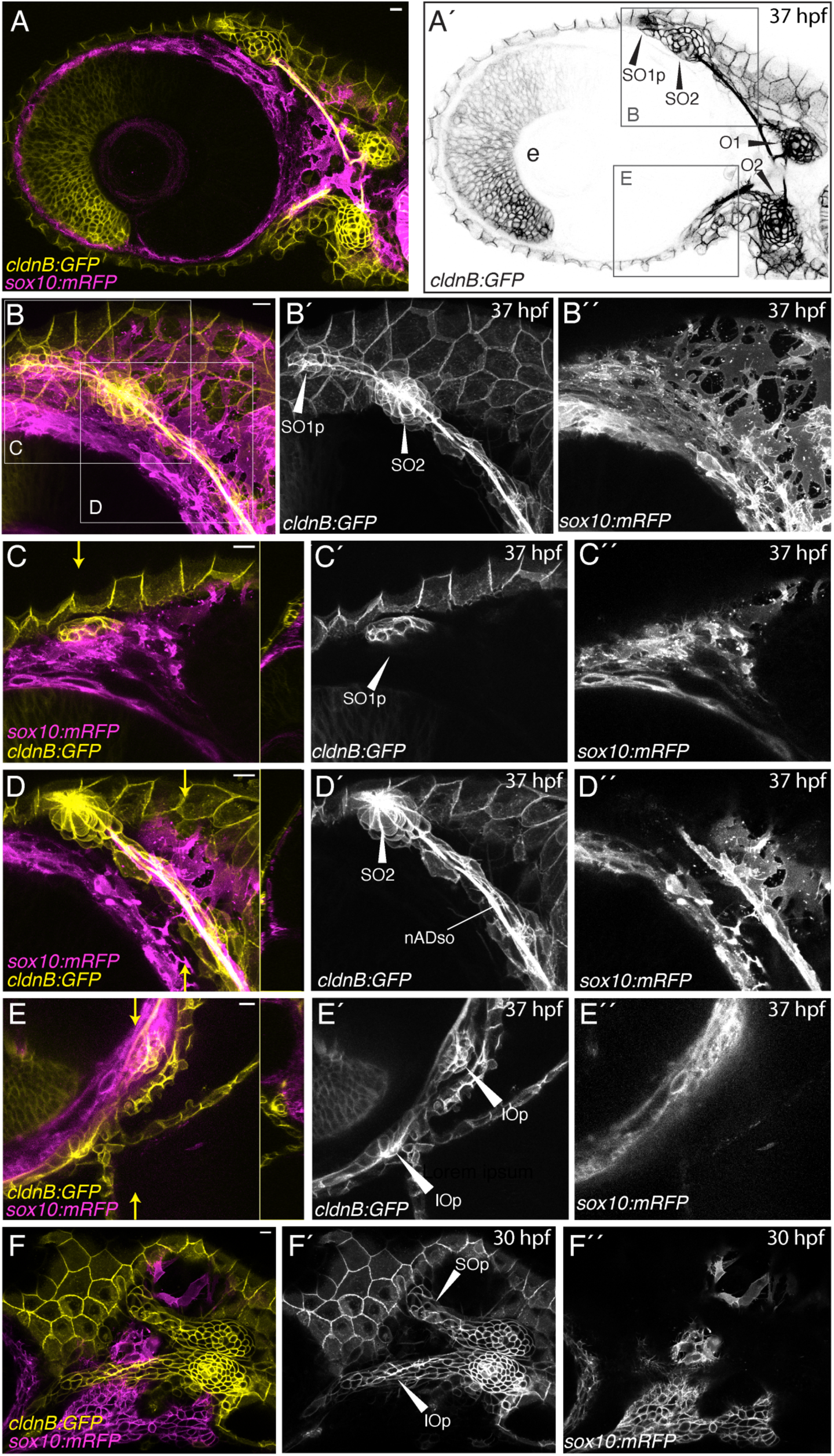
Confocal imaging reveals that cells of the anterior lateral line develop in close proximity to cranial neural crest cells. (**A**) Lateral view of the cranial region of a 37 hpf *Tg(cldnB:GFP;sox10:mRFP)* double transgenic embryo, anterior to the left. Maximum projection of a 3 µm deep z-stack, imaged using the 10x objective, shows the juxtaposition of the developing anterior lateral line (yellow) with cranial neural crest cells (magenta). (**A’**) Single channel image (*cldnB:GFP*) of the same specimen, showing the developing lateral line system only. Neuromasts are labeled:-SO2: supraorbital neuromast 2; SO1p: primordium of supraorbital neuromast 1 in the process of budding from SO2; O1, O2: otic neuromasts; e: lens of the eye. Boxed regions indicate the approximate regions imaged in panels B and E. (**B-E**) Maximum projections of a second 37 hpf *Tg(cldnB:GFP;sox10:mRFP)* specimen, imaged using the 40x objective. (**B-B’’**) Maximum projection of a 30.75 µm deep z-stack, centered on the developing supraorbital region (see panel A’ for region). Hatched boxes indicate regions imaged at higher magnification in panels C and D. (**C-C’’**) Maximum projection of a 3.75 µm deep sub-stack of the z-series from the specimen in B, in a focal plane showing the SO1 primordium (SO1p, indicated in C’; *cldnB:GFP*) budding over neural crest cells (C’’; *sox10:mRFP*). An orthogonal yz reslice through the whole stack, in the plane indicated by small yellow arrows, is provided to the right of C. (**D-D’’**) Maximum projection of a 3.75 µm deep sub-stack of the z-series from the specimen in B, in a focal plane showing the SO2 neuromast and supraorbital nerve (nADso; D’). An orthogonal yz reslice through the whole stack, in the plane indicated by small yellow arrows, is provided to the right of D. Note the nSO nerve is wrapped in neural crest cells (D’’; *sox10:mRFP*). (**E-E’’**) Maximum projection of a 3.75 µm sub-stack of the larger z-series, in a focal plane showing the developing infraorbital region (see panel A’ for region). An orthogonal yz reslice through the whole stack, in the plane indicated by small yellow arrows, is provided to the right of E. IOp indicates the infraorbital primordium (E’; *cldnB:GFP*) elongating over neural crest cells (E”; *sox10:mRFP*). (**F-F’’**) A single (1.44 µm) z-slice of a 30 hpf *Tg(cldnB:GFP;Sox10:mRFP)* embryo, imaged using the 40x objective, showing supra- and infraorbital primordia (SOp and IOp) of the anterior lateral line diverging away from one another (F’) over a field of neural crest cells (F”; magenta). Scale bars are 10 µm throughout.

Fig. 2C is a maximum projection of three z-slices (total z depth = 3.75 µm) in the plane of the SO1 primordium, revealing that the *cldnB:GFP*-expressing lateral line cells budding from SO2 (Fig. 2C’; SO1p) overlie *sox10:mRFP-*expressing neural crest cells (Fig. 2C”). An orthogonal yz reslice through the whole z-stack is shown on the right-hand side of Fig. 2C, confirming the close proximity of the two cell types. Fig. 2D is a maximum projection of three z-slices (total z depth = 3.75 µm) in the plane of SO2 and the supraorbital nerve (Fig. 2D’, nADso), revealing that *sox10:mRFP-*expressing neural crest cells wrap around the *cldnB:GFP*-expressing SO nerve (compare Figs. 2D, D’, D”; an orthogonal yz reslice through the whole z-stack is provided on the right-hand side of Fig. 2D). Fig. 2E shows a maximum projection of three z-slices in the infraorbital region (total z depth = 3.75 µm; an orthogonal yz reslice through the whole z-stack is provided on the right-hand side). The infraorbital, like the supraorbital, has *cldnB:GFP*-expressing lateral line cells (Fig. 2E’) in close contact with underlying *sox10:mRFP-*expressing neural crest cells (Fig. 2E”), confirming the close proximity of the two cell types. Finally, to address whether the close relationship between placode-derived and neural crest cells is present earlier in development, we performed a similar analysis at the 30 hpf stage, when the SO and IO primordia are first forming. Fig. 2F is a single z-slice (z depth 1.44 µm) that shows both the SO and IO primordia branching away from one another. Already at this stage, the *cldnB:GFP*-expressing lateral line cells (Fig. 2F’) are directly overlying *sox10:mRFP-*expressing neural crest cells (Fig. 2F”).

Together, our lightsheet and confocal microscopy analyses confirm that placode-derived anterior lateral line primordia develop in close proximity to cranial neural crest cells, from as early as 30 hpf through the first 5 days of larval development.

### 2.2 Lateral line ganglia have a shared neural crest and placode origin

The lateral line system not only comprises the neuromast sensory organs, but also the ganglia and afferent nerves that innervate them. Moreover, all these components of the system derive from the same set of cranial placodes. We therefore turned our attention to investigating the anterior lateral line ganglia and their relationship with the neural crest cells.

Figure 3A-D’’ shows lateral views of confocal maximum projections of the cranial region of *Tg(cldnB:GFP)* (yellow) specimens at 24 hpf, 30 hpf, 36 hpf, and 48 hpf. Specimens were immunolabeled with anti HuC/D antibody, a marker of differentiated neurons (cyan), and anti-Sox10 antibody, which marks neural crest cell nuclei (magenta). The anterodorsal component (gAD) of the ALL ganglion is specified by 24 hpf (Fig. 3A-A’’), adjacent to the acoustic ganglion (gVIII), and shifts anteriorly towards the trigeminal ganglion (gV) by 30 hpf, while rapidly growing in size (Fig. 3B-B’’), as previously reported (Raible & Kruse, 2000). At 36 hpf and 48 hpf, the gAD retains a spherical shape and lies directly adjacent and posterior to the trigeminal ganglion (Fig. 3C-3D’’). We were able to observe the first signs of neurons in the anteroventral component (gAV), specified ventromedially to the gAD, at 36 hpf (Fig. 3C-C’’), a few hours earlier than the previously reported 40 hpf (Raible and Kruse 2000). Neurons of the lateral line ganglia (gAD, gAV, gP) can be identified by double labeling with HuC/D (cyan) and CldnB:GFP (yellow) (Fig 3A’’-D’’). Throughout the development of these lateral line ganglia, Sox10-positive neural crest cell nuclei are clustered around the gAD, gAV, and gP (Fig. 3A-D). We include a single z-slice inset (z depth 1.22 µm) in Fig. C’’, which shows how neural crest cells (magenta) directly contact the clustered HuC/D-positive, CldnB:GFP-positive (cyan and yellow, respectively) cells of the gAD. We also observed that neural crest cells wrap around the supraorbital (nADso) and infraorbital (nADio) lateral line nerves of the gAD as early as 30 hpf (Fig. 3B’’-D’’), as well as the mandibular nerve (nAVmd) of the gAV once it is specified (Fig. 3D’’’).

**Figure 3:**
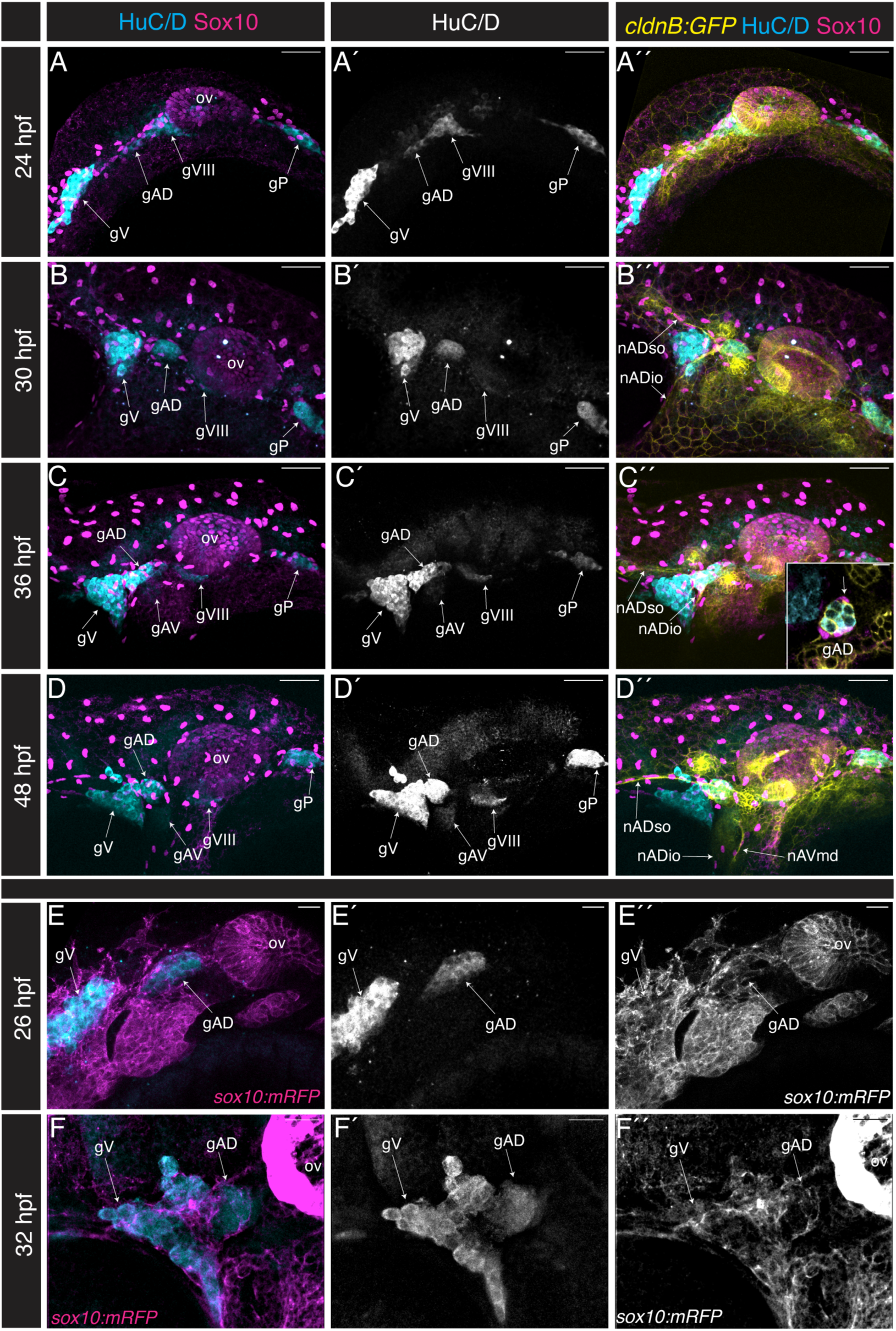
Lateral line ganglia form in close association with neural crest cells. (**A-D’’**) Confocal maximum projections in lateral view (40x objective), anterior to the left, of the lateral line ganglia of specimens co-labeled for differentiated neurons (anti HuC/D; cyan), neural crest cells (Sox10; magenta) and lateral line (*Tg(cldnB:GFP):* yellow); channels as indicated. In A (24 hpf), B (30 hpf), C (36 hpf), and D (48 hpf) neural crest cell nuclei are labeled by anti-Sox10 antibodies. Inset in C’’ is a single z-slice (z depth = 1.22 µm) showing neural crest cells (magenta) enveloping neurons in the anterodorsal lateral line ganglion (arrow; gAD; cyan). (**E-F’’**) In E (26 hpf) and F (32 hpf) neural crest cell membranes are labeled by *Tg(sox10:mRFP)* (magenta) and differentiated neurons are labeled with anti HuC/D (cyan). The anti-HuC/D marker is shown in single channel in E’ and F’; *Tg(sox10:mRFP)* is shown in single channel in E” and F”. Abbreviations:-ov: otic vesicle; gV: trigeminal (Vth nerve) ganglion; gAD: anterodorsal lateral line ganglion; gAV: anteroventral lateral line ganglion; gVIII: acoustic ganglion; gPLL: posterior lateral line ganglion; nADso: supraorbital nerve; nADio: infraorbital nerve; nAVmd: mandibular nerve.

Expression of Sox10 protein is downregulated rapidly in some migrating neural crest cells, such that anti-Sox10 antibody does not detect neural crest cell nuclei in those cells populating the developing pharyngeal arches (Fig. 3A-D’’). By contrast, the membrane RFP signal provided by *Tg(sox10:mRFP)* is long-lasting, providing a more complete readout of neural crest cell localization. We therefore immunolabeled *Tg(sox10:mRFP)* specimens with anti-HuC/D antibody, allowing a comparison of neural crest cell and neuron localization at 26 and 32 hpf (Fig. 3E-F’’). At both stages, we find that the gAD is encased in a ‘shell’ of neural crest tissue.

While data in Figure 3 indicate that lateral line gangliogenesis occurs in close proximity to the neural crest, it is unclear from this analysis whether neurons within the ganglia are neural crest cell-derived. To investigate the neural crest contribution to the anterior lateral line ganglion in more detail, we utilized Cre-based lineage tracing of the neural crest cells, using a two-transgene system previously employed by Kague et al. (2012): *Tg(−28.5Sox10:cre;ef1a:loxP-dsRed-loxP-egfp)*, for simplicity referred to hereafter as *Tg(sox10Cre;dsRed/EGFP*). The first transgene uses the neural crest-specific *sox10* regulatory sequences to drive expression of Cre recombinase, and the second transgene responds as a “switch”, with LoxP sites recombined by Cre to remove dsRed gene and bring EGFP under control of the EF1alpha ubiquitous promoter. Thus, despite the down-regulation of Sox10 expression described above, once cells have expressed the *sox10*-driven Cre recombinase, they become genetically labeled by EGFP, a trait that is passed on to all that cell’s progeny.

Figure 4A, B shows a lateral view of a confocal projection of the cranial region of a 120 hpf *Tg(sox10Cre; dsRed/EGFP*) specimen, with the organization of the cranial ganglia again revealed by immunolabeling with the HuC/D marker of differentiated neurons (cyan). Fig. 4B shows the location of EGFP-labeled neural crest cells (magenta). Fig. 4C is a schematized version of Fig. 4B, indicating the locations of cranial ganglia and summarizing their placodal (cyan) or placode plus neural crest (hatched cyan/magenta) origins. We find, confirming the previous report by Kague et al. (2012), that the anterior lateral line, posterior lateral line, Vth/trigeminal, VIIth/facial, and VIIIth/acoustic ganglia (Fig. 4C: gP, gV, gVII, and gVIII) each include neurons of mixed origins. In Figs 4D, E, we extend these previous findings, to demonstrate that both the dorsal and the ventral components of the anterior lateral line ganglion (gAD and gAV) share placode and neural crest cell contributions. Figs 4D, E are single high magnification confocal slices through the adjoining trigeminal (gV) and anterior lateral line ganglia (region boxed with red dashes in Fig. 4C). Panels D-D’ show a single confocal z-slice through the superficial dorsal ganglion (gAD), whereas panels E-E’ show a single confocal z-slice through the deeper ventral ganglion (gAV). Fig. 4F schematizes the relative locations of these slices. All neurons are HuC/D-positive (Figs 4D, D’’ and E, E’’) and a subset of the gV, gAD and gAV ganglia are EGFP-labeled (magenta) (Figs 4D’, E’), revealing their neural crest origins. In summary, both the anterodorsal and the later forming anteroventral components of the zebrafish anterior lateral line ganglion combine placode-derived and neural crest-derived neurons (schematized in Fig. 4G).

**Figure 4:**
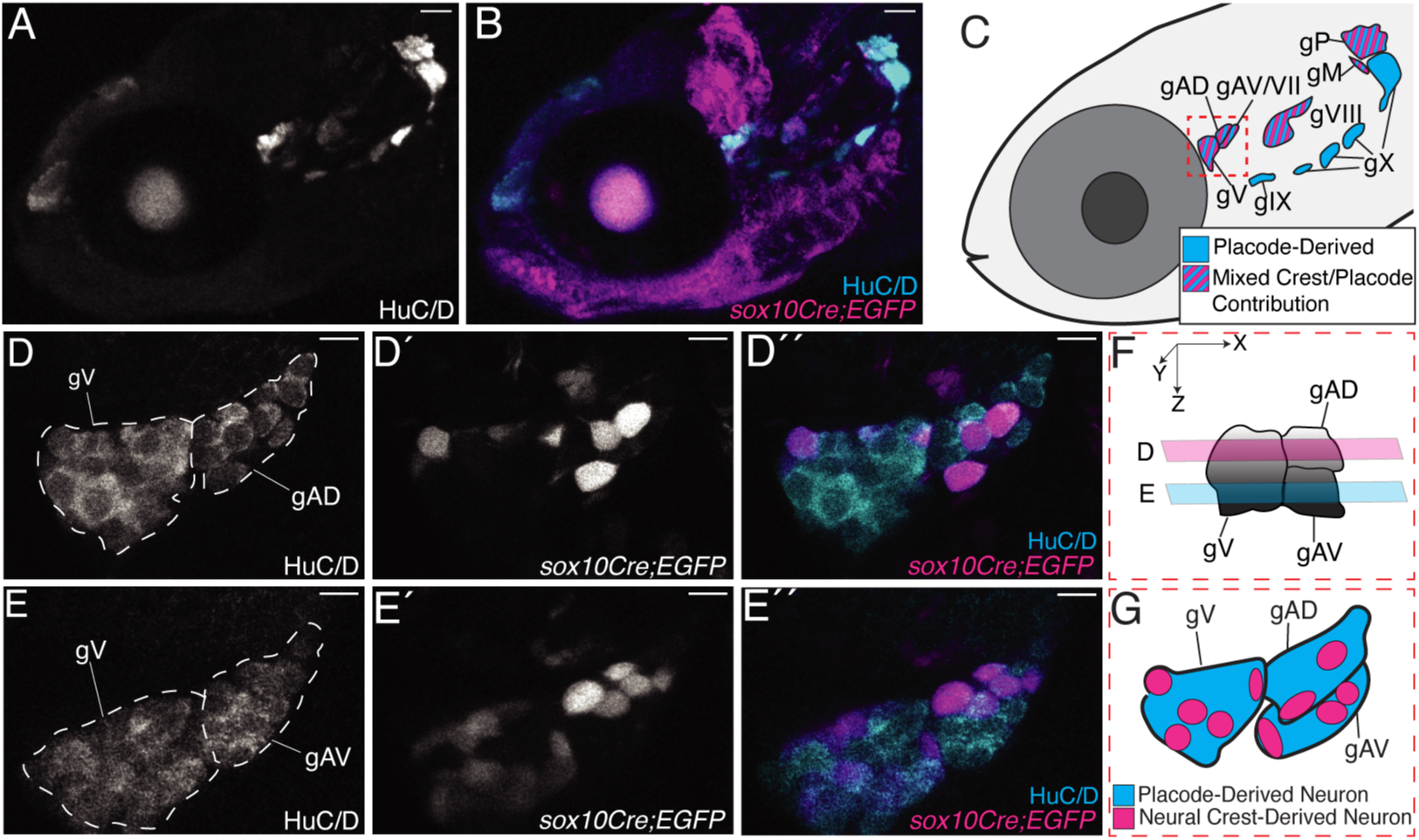
Lateral line ganglia have a shared neural crest and placode origin. (**A-C**) Confocal maximum projection (10x objective), in lateral view with anterior to the left, of the cranial region of a *Tg(sox10Cre;dsRed/EGFP*) 120 hpf specimen immunolabeled with anti HuC/D (of n=6). (A) HuC/D-positive differentiated neurons only. (B) HuC/D-positive neurons (cyan) together with *sox10Cre;EGFP* neural-crest derived cells (magenta). (C) Schematic representation of panel B indicating placode and neural crest contributions to the cranial ganglia. Abbreviations are as follows:-gV: trigeminal ganglion; gAD anterodorsal lateral line ganglion; gAV/gVII: anteroventral lateral line ganglion fused with facial ganglion; gVIII: acoustic ganglion; gM: middle lateral line ganglion; gP: posterior lateral line ganglion, gIX: glossopharyngeal ganglion; gX: vagal ganglion. (**D-E’’**) Higher magnification views (40x objective) of the region boxed in C. (**D-D’’**) A single z-slice (1.02 µM), in the plane of the anterodorsal lateral line ganglion and trigeminal ganglion. (D) HuC/D-positive neurons; (D’) neural-crest-derived cells (*Sox10Cre;EGFP*); (D’’) merge, neural-crest derived cells in magenta and HuC/D-positive neurons in cyan. (**E-E’’**) A single z-slice (1.02 µM), in the plane of the anteroventral lateral line and trigeminal ganglion. (E) HuC/D-positive neurons; (E’) neural crest-derived cells (*Sox10Cre;EGFP*); (E’’) merge, neural-crest derived cells in magenta and HuC/D-positive neurons in cyan. (**F**) Schematic representation of the ganglia shown in D and E, indicating the z-planes imaged. (**G**) Schematic representation of panels D’’ and E’’, showing the neural crest contribution (magenta) to the anterior lateral line ganglia and the trigeminal ganglion. Scale bars are 50 µm in A-C, and 10 µm in D-E.

### 2.3 Anterior lateral line development is disrupted in the absence of neural crest

Having confirmed a close physical relationship between cranial neural crest cells and all parts of the anterior lateral line system, we tested the hypothesis that neural crest cells play a functional role in anterior lateral line patterning and development. To remove neural crest cells we turned to a commonly used method to ablate zebrafish cells conditionally: the genetic/pharmacological approach based on expression of the transgene-encoded bacterial enzyme Nitroreductase (NTR) (Curado et al., 2007). Genetically encoded NTR converts a prodrug into a cytotoxic compound that is strictly limited to the NTR-expressing cells, thus causing cell-specific apoptosis (White & Mumm, 2013). To achieve neural crest-specific cell death we used the *Tg(sox10:NTRmCherry)* transgene, which expresses NTR enzyme and mCherry fluorescent protein under control of the neural crest specific *sox10* regulatory sequences. By treating these transgenic specimens with the prodrug Nifurpirinol (NIF) we achieved effective ablation of the NTR-expressing neural crest cells without generalized toxicity, consistent with previous reports (Bergemann et al., 2018; Cavanaugh et al., 2015). We then used this approach to evaluate how depletion of neural crest cells impacts development of the anterior lateral line system.

Figure 5A-D’ shows lateral views of confocal maximum projections of the crania of *Tg(sox10:NTRmCherry; cldnB:GFP)* double transgenic specimens, treated with DMSO carrier control, at 36 hpf, 48 hpf, 72 hpf, and 120 hpf. As expected, development of the lateral line system in the DMSO-treated controls was indistinguishable from that of unmanipulated specimens (compare Fig. 5A-B’ with Figs 1 and 2). Fig. 5E-L shows *Tg(sox10:NTRmCherry;cldnB:GFP)* experimental specimens, which were treated with 1.25 µM NIF from the 9 hpf stage; a stage selected because it is immediately before the onset of *sox10* expression. In the experimental specimens, at 36 hpf and at all subsequent stages, we observed widespread neural crest cell death: apoptotic cells are distinguished by their reduced size and eventual degradation to puncta, accompanied by a dramatic reduction in mCherry signal. The absence of neural crest cells led to disrupted cranial morphology, with a complete absence of jaw structures apparent by 72 hpf. Despite a lack of neural crest cells the specimens remained generally healthy and continued to develop, although by 120 hpf they displayed dysmorphic hearts as previously reported in response to an equivalent manipulation (Cavanaugh et al., 2015). We also noted that the otic vesicle was reduced in size, consistent with expression of *Tg(sox10:NTRmCherry)* in this structure. We used the *cldnB:GFP* transgene to evaluate how the lateral line system was affected by the loss of neural crest cells. Fig. 5E-L shows two independent experimental specimens out of n=/>10, at each of the four developmental stages.

**Figure 5:**
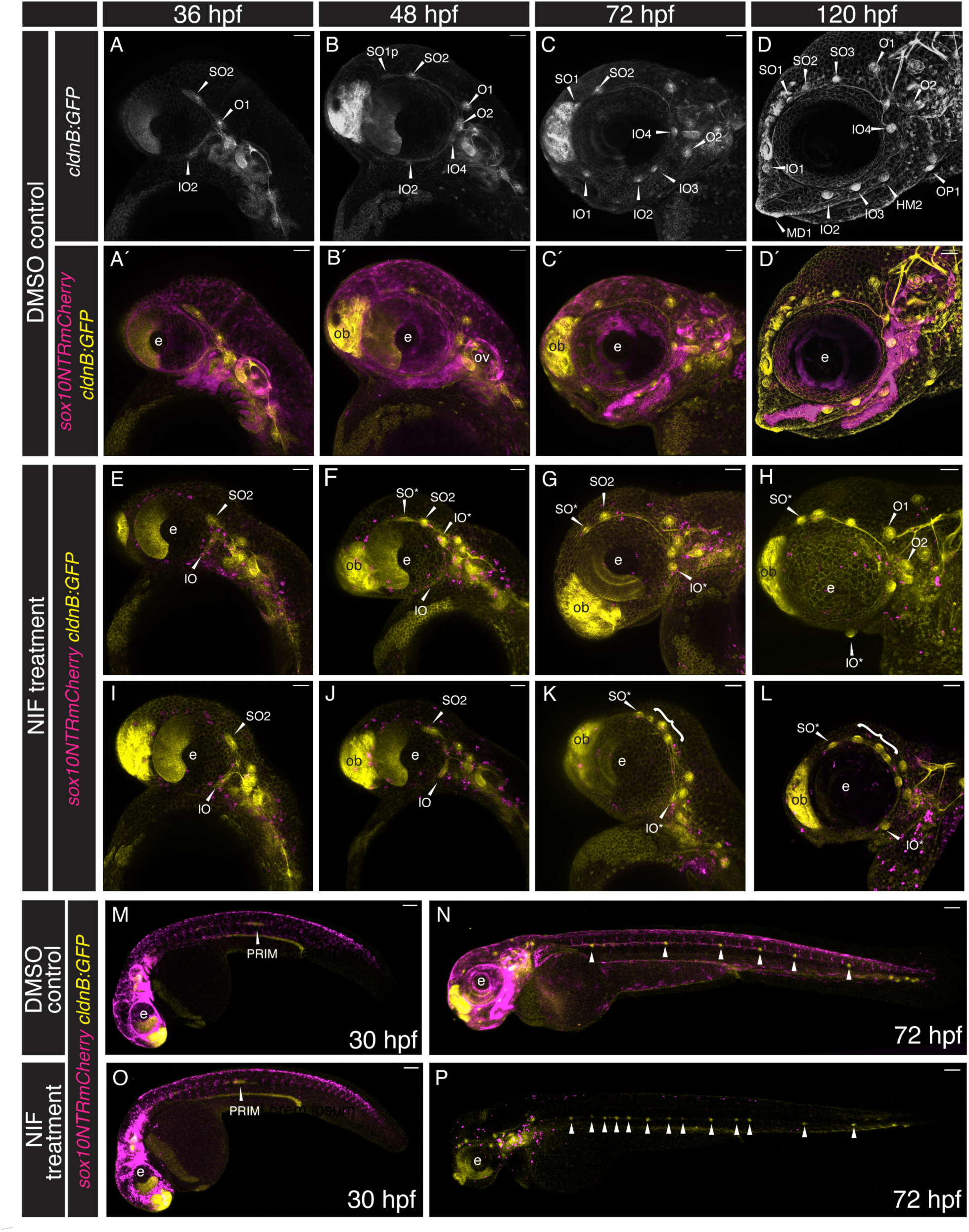
Anterior lateral line development is disrupted in the absence of neural crest cells. (**A-L**) Confocal maximum projections, in lateral view with anterior to the left, of the cranial regions of *Tg(sox10:NTRmCherry;cldnB:GFP)* double transgenic specimens at 36 to 120 hpf. (**A-D’**): DMSO carrier-treated controls show normal development of the lateral line; stages are as indicated. A-D show the lateral line alone *(cldnB:EGFP)*; A’-D’ show both lateral line (*cldnB:EGFP*; yellow) and neural crest (*sox10:NTRmCherry*; magenta). (**E-L**) When neural crest (*sox10:NTRmCherry*; magenta) cytotoxicity is activated via Nifurpirinol (NIF) treatment, lateral line (*cldnB:GFP*; yellow) development is disrupted. Two different samples are shown at each stage (n= 14 at 36 hpf, n=10 at 48 hpf, n= 10 at 72 hpf, n= 16 at 120 hpf). The dying *sox10:NTRmCherry*-positive neural crest cells show punctate fluorescence at all stages, with the number of remaining neural crest cells reducing over time. Lack of neural crest leads to disrupted morphogenesis of the anterior lateral lines. The anterior-most neuromast of the supraorbital line (SO*) never reaches the olfactory bulb (ob), even at 120 hours post fertilization. By 72 hpf we often observe an array of ectopic neuromasts in the posterior supraorbital domain (brackets, K, L). Infraorbital lateral line development is severely truncated, with fewer than normal infraorbital neuromasts, often inappropriately positioned (IO*), developing. (**M-P**) Low magnification, tiled, confocal maximum projections, in lateral view with anterior to the left, of entire *Tg(sox10:NTRmCherry;cldnB:GFP)* double transgenic specimens. (**M, N**): DMSO carrier-treated control specimens show normal development of the posterior lateral line (*cldnB:EGFP*; yellow) and trunk neural crest cells (*sox10:NTRmCherry*; magenta). (M) 30 hpf, the posterior lateral line primordium (PRIM; yellow) is migrating through the anterior trunk, and neural crest cells (magenta) are located in the dorsal neural tube and migrating ventrally in segmental streams. (N) 72 hpf, shows six evenly distributed trunk neuromasts (arrowheads). (**O, P**) Nifurpirinol (NIF) treatment causes neural crest (*sox10:NTRmCherry*; magenta) cytotoxicity and formation of supernumerary trunk neuromasts, but does not prevent the posterior lateral line from migrating fully along the trunk. (O) 30 hpf; shows the posterior lateral line primordium (PRIM; yellow) migrating normally through the anterior trunk, while neural crest cells (magenta) are starting to die. (P) 72 hpf; shows multiple supernumerary trunk neuromasts (arrowheads), while most neural crest cells (magenta) are now absent. Abbreviations are as follows:-e: lens of the eye; ov: otic vesicle; ob: olfactory bulb; SO: supraorbital neuromast; IO: infraorbital neuromast; O: otic neuromast; OP: opercular neuromast; HM: hyomandibular neuromast; MD: mandibular neuromast. Scale bars are 50 µM throughout.

At the 36 hpf stage there were only modest differences between the anterior lateral lines of control (Fig. 5A, A’) and NIF-treated experimental (Fig. 5E, I) specimens. When neural crest cells were depleted, the supraorbital primordium exhibited a modest reduction in distance migrated above the eye, but otherwise appeared similar to control specimens. The infraorbital cell stream similarly exhibited a more limited extension relative to controls, and also appeared slightly broader. By the 48 hpf stage the phenotypes in experimental specimens were more pronounced. In 48 hpf control specimens (Fig. 5B, B’) the SO1 neuromast had already begun to bud from SO2, and extended anteriorly, approaching the developing olfactory bulb (Fig. 5B’, ob). By contrast, in the experimental specimens (Fig. 5F, J) the supraorbital line had not extended beyond the dorsal apex of the eye, with limited or scant SO2 budding (Fig. 5F, SO*; Fig, 5J). Additionally in 48 hpf control specimens, the infraorbital line had completely extended along the ventral edge of the eye (Fig. 5B, B’), and the IO2 neuromast had formed, whereas in experimental specimens the infraorbital was significantly underdeveloped, and in most cases had extended very little beyond its 36 hpf position with no IO2 neuromast in place (Fig. 5F, J). In some experimental specimens an infraorbital neuromast appeared to have been displaced dorsally, up into the supraorbital region (e.g. Fig. 5F, IO*). Additional evidence supporting the assignment of such ectopic neuromasts to the infraorbital system came from our studies of innervation and time-lapse microscopy (see Figs 6-8, below).

**Figure 6:**
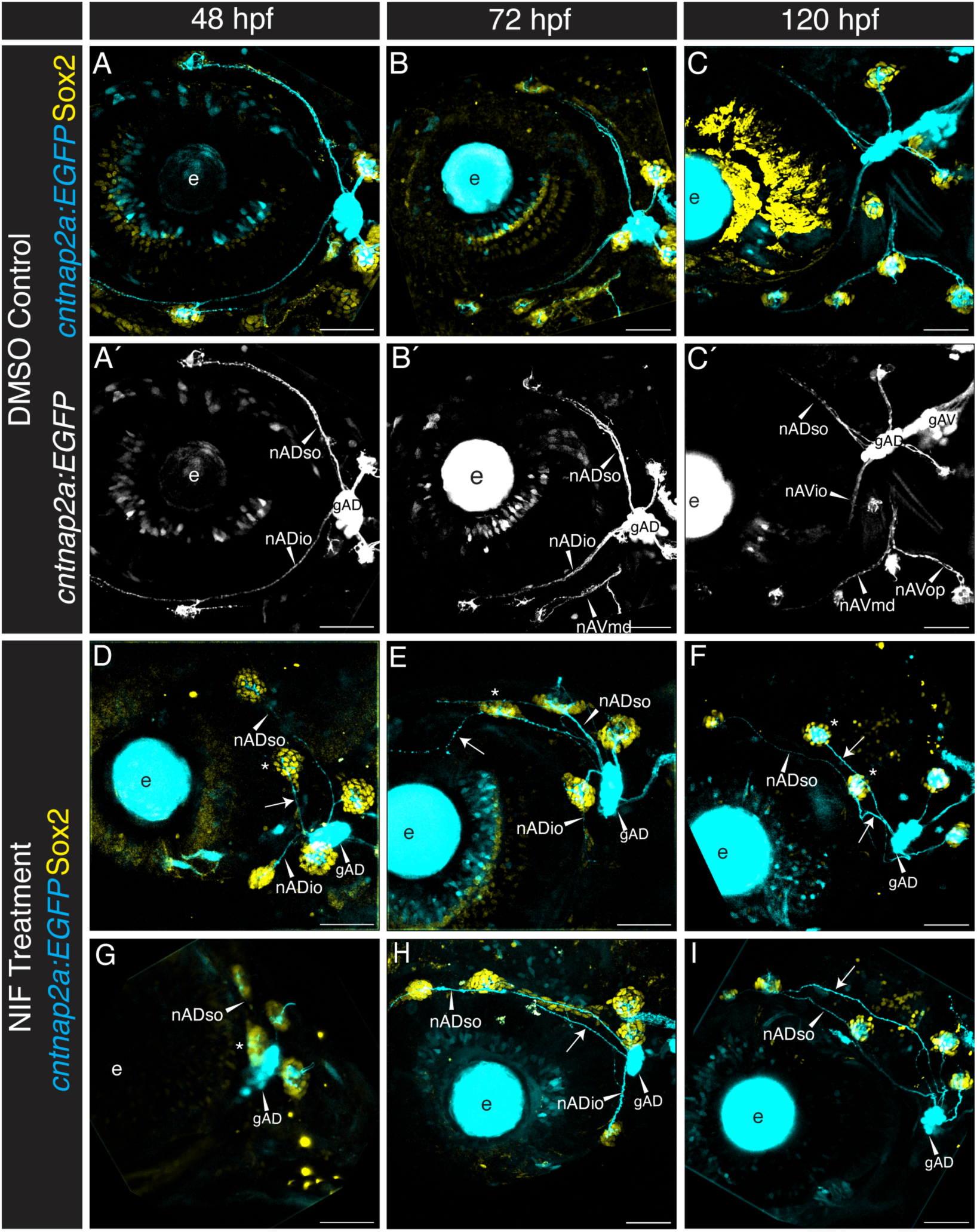
Lateral line innervation is disrupted in the absence of neural crest. (**A-I**) Confocal maximum projections (40xobjective), in lateral view with anterior to the left, of the cranial regions of *Tg(sox10:NTRmCherry;cntnap2a:EGFP)* double transgenic specimens at 48 to 120 hpf. The *cntnap2a:EGFP* transgene (cyan) labels afferent lateral line neurons and the lens of the eye. The specimens are also immunolabeled with the neuromast support cell marker anti-Sox2 (yellow). (**A-C**): DMSO carrier-treated controls show normal development of the lateral line nerves (*cntnap2a:EGFP*; cyan) and neuromasts (Sox2; yellow). (**A’-C’**) are the same specimens as A-D, shown in the *cntnap2a:EGFP* (cyan) channel alone. Stages are as indicated. (**D-I**) When neural crest (*sox10:NTRmCherry*; not shown) cytotoxicity is activated via Nifurpirinol (NIF) treatment, lateral line innervation (*cntnap2a:EGFP*; cyan) and neuromast organization (Sox2, yellow), are both disrupted. Two different samples, out of n=10 at each stage, are shown. Innervation is disrupted, with gAV nerves nAVmd and nAVop failing to form by 120 hpf (compare F, I with C’). Nerves often project to misplaced neuromasts, although some fail to reach a neuromast and project ectopically (arrows). Nerves also make multiple supraorbital projections, often innervating ectopic or supernumerary neuromasts (D, E, F, G, asterisks), and tend to be defasiculated and contain fewer axons in comparison to DMSO-treated control specimens. Abbreviations are as follows:-e: lens of the eye; nADso: supraorbital nerve; nADio: infraorbital nerve; nAVmd mandibular nerve; nAVop: opercular nerve. Scale bars are 50 µM throughout.

The phenotype of restricted supra and infraorbital development continued through the 72 and 120 hpf stages in NIF-treated specimens (Fig. 5G, H, K, L). The supraorbital line rarely extended beyond the apex of the eye, never reaching the olfactory bulb region, while the infraorbital line remained drastically shortened with missing neuromasts (compare Fig. 5C, C’ with Fig. 5G, K, and Fig. 5D, D’ with Fig. 5H, L). At the 120 hpf stage, some experimental specimens had formed an additional infraorbital neuromast, to result in two total (compare Fig. 5H, L with D, D’). We never observed more than three total IO neuromasts in experimental specimens, whereas control specimens always had four IO neuromasts (Fig. 5D, D’). Additionally, at these later stages, we frequently observed emergence of ectopic supernumerary neuromasts in the posterior supraorbital region (brackets, Fig. 5K, L), resulting in an array of closely clustered neuromasts. By 120 hpf the mandibular and opercular lines have begun to form in control specimens (Fig. 5D, D’). Both are entirely absent from the neural crest deficient specimens, consistent with major deficits in the neural crest-populated mandibular and hyoid arches.

In the absence of neural crest-derived glial cells, as occurs in several zebrafish mutants including those in the ErbB/Neuregulin pathway (Lush & Piotrowski, 2014), precocious intercalary neuromasts form in the posterior lateral line system. We found that our neural crest cell-deficient specimens similarly displayed supernumerary posterior lateral line neuromasts in the trunk, consistent with the predicted loss of glial cells. Figure 5M-P shows confocal projections of whole *Tg(sox10:NTRmCherry;cldnB:GFP)* larvae in lateral view at the 30 hpf and 72 hpf stages. DMSO-treated control specimens display typical posterior lateral line primordium migration at 30 hpf, with healthy neural crest cells (magenta) migrating ventrally in streams adjacent to each somite (Fig. 5M). By 72 hpf, 5-7 primary posterior lateral line neuromasts have been deposited at regular intervals along the trunk (Fig. 5N, this example has 6 trunk neuromasts). At 30 hpf the NIF-treated specimens showed normal primordium migration, although the most anterior neural crest cells—which were born earliest—are already rounding up and undergoing apoptosis (Fig. 5O). By 72 hpf all neural crest cells are either absent or dead, and supernumerary posterior lateral line neuromasts are present (Fig. 5P, this example has 13 trunk neuromasts). Importantly, despite the development of precocious neuromasts, the posterior lateral line migrates all the way to the posterior limit of the trunk when neural crest cells are absent (n=10). We conclude that migration of the posterior lateral line primordium is independent of neural crest cells, unlike the situation for the anterior lateral line system.

### 2.5 Anterior lateral line gangliogenesis and innervation are disrupted in the absence of neural crest

As cranial neural crest cells contribute directly to the lateral line ganglia (Figs 3, 4) we predicted that absence of neural crest would cause disruptions to anterior lateral line gangliogenesis and potentially to innervation. We tested this prediction by once again taking advantage of *Tg(sox10:NTRmCherry)* specimens and NIF treatment to cause neural crest cell death. Figure 6A-C shows lateral views of confocal projections of the region surrounding, and posterior to, the developing eye of DMSO control-treated *Tg(sox10:NTRmCherry)* specimens at 48 hpf, 72 hpf, and 120 hpf. These specimens additionally carry *Tg(cntnap2a:EGFP)* (cyan), which is expressed exclusively in the afferent lateral line nerves (Faucherre et al., 2009; Pujol-Martí et al., 2012), and are immunolabeled with anti-Sox2 antibody (yellow), to mark neuromast support cells (Hernández et al., 2007); Figs 6A’-C’ show *cntnap2a:EGFP* alone. At the 48 hpf stage (Fig. 6A, A’) the dorsally located SO2 neuromast is innervated by the supraorbital nerve (nADso) and the ventrally located IO2 neuromast is innervated by the infraorbital nerve (nADio). Both the supraorbital and infraorbital nerves extend from the gAD ganglion. Two additional nerves emerge from gAD to innervate the otic neuromasts, O1 and O2, generating a characteristic ‘X’ pattern, which is maintained at 72 hpf and 120 hpf as development proceeds (Fig. 6B, B’; C, C’). By 72 hpf (Fig. 6B, B’) the developing mandibular neuromasts are innervated by the mandibular nerve (nAVmd), which emerges from the recently formed ventral (gAV) anterior lateral line ganglion. By 120 hpf this nerve has branched to form the opercular nerve (nAVop), and the ventral ganglion (gAV) can now be visualized as a separate component of the anterior lateral line ganglion.

The DMSO-treated control specimens (Fig. 6A-C’) are compared with two independent NIF-treated specimens (out of n=10 at each stage; Fig. 6D-I). As early as 48 hpf, ectopic nerves could be visualized in the supraorbital region (arrows), frequently innervating misplaced neuromasts (indicated by asterisks; e.g. Fig. 6D, E, F). The dorsal translocation of the nADio nerve confirmed that an IO neuromast is often deposited in an unusually dorsal location to lie close to SO neuromasts (e.g. Fig. 6E). By 120 hpf, 8 of 10 NIF-treated specimens have multiple dorsally-directed projections (e.g. Fig. 6F, I; and see also Fig. 7F, ahead), suggesting that ectopic supernumerary neuromasts originate not only from the misdirected IO, but additionally from other sources, which might include precocious proliferation of interneuromast cells. By 72 hpf the gAD ganglion of NIF-treated specimens is reduced in size relative to DMSO-treated controls, and by 120 hpf it is apparent that the gAV ganglion is missing, as are the mandibular and opercular nerves (Fig. 6F, I). In addition to reduced and missing ganglia, the nerves themselves are often thinner in the NIF-treated neural crest-deficient specimens, likely as a consequence of missing neural crest-derived neurons. We also noted that the nerves emerged from non-uniform points on the ganglia, relative to controls, and that the nerves are often defasciculated, as additionally shown ahead (see Fig. 7D). In some specimens, nerves extended erratically in the supraorbital region without innervating a neuromast (e.g. Fig. 6E).

**Figure 7:**
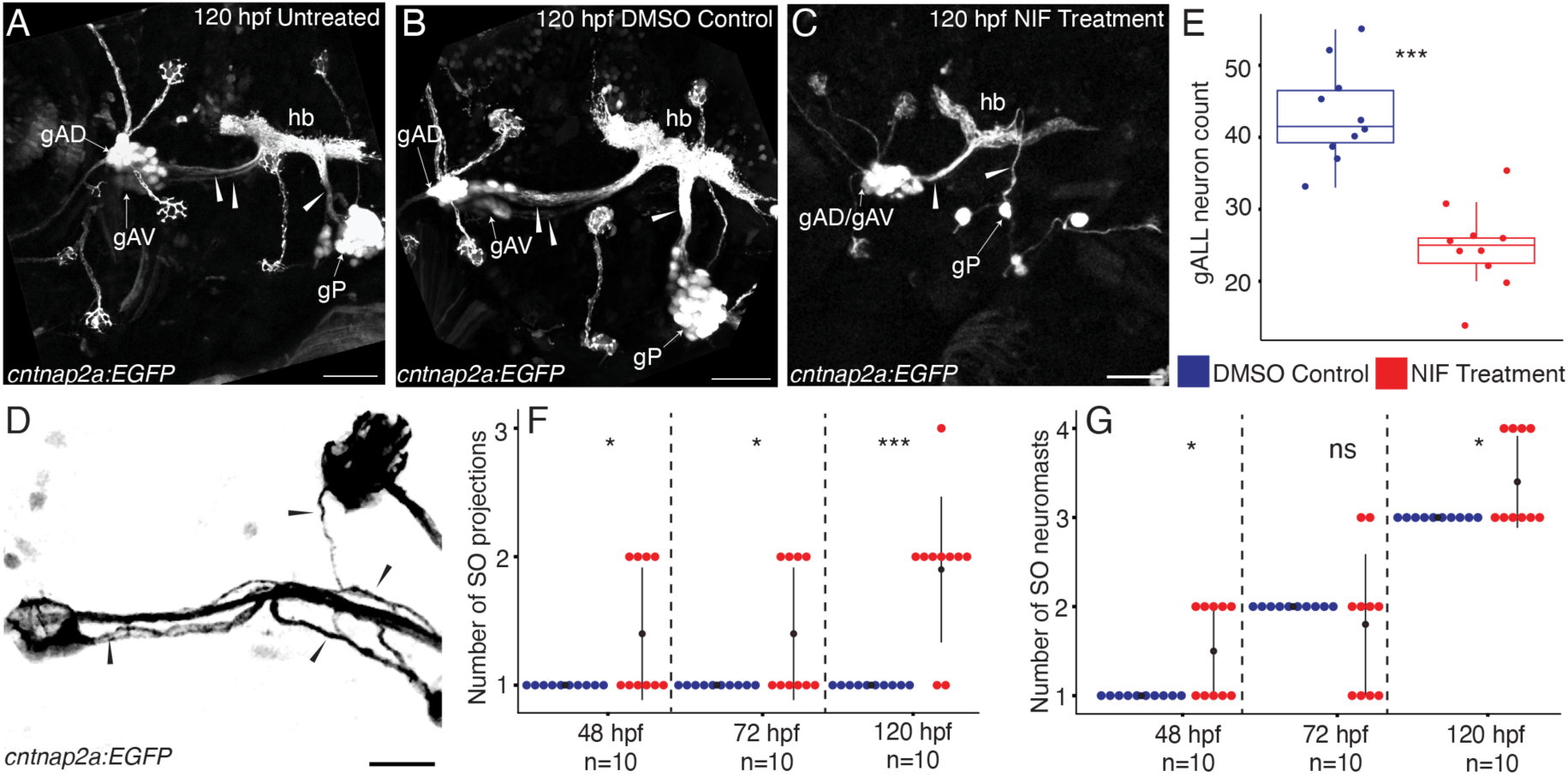
Lateral line patterning and gangliogenesis are disrupted in the absence of neural crest. (**A-C**) Confocal maximum projections (40x objective), in lateral view with anterior to the left, of the lateral line afferent system of *Tg(sox10:NTRmCherry;cntnap2a:EGFP)* specimens at 120 hpf; *(cntnap2a:EGFP)* shown in single channel. (A) Untreated specimen showing the locations of the gAD, gAV, and gP lateral line ganglia and their individual projections to the hindbrain (arrowheads). (B) DMSO carrier-treated control showing normal ganglia morphology and innervation patterns *(cntnap2a:EGFP)*. (C) When neural crest (*sox10:NTRmCherry*) cytotoxicity is activated via Nifurpirinol (NIF) treatment, lateral line ganglion morphology is disrupted. The gAV and gAD ganglia are no longer separated, show altered density and shape, and make a shared projection to the hindbrain. The posterior ganglion has severe deficits. (**D**) Example of neuromast-innervating nerves (*cntnap2a:EGFP*, black) defasciculation (arrowheads) in the absence of neural crest cells. (**E**) Boxplot showing counts of HuC/D-expressing neurons, in both the dorsal and ventral parts of the anterior lateral line ganglia in DMSO control versus NIF-treated specimens at 120 hpf (n=10 for each condition). (**F**) Stacked Histogram showing the number of supraorbital nerve projections to neuromasts in DMSO control versus NIF-treated conditions at 48 hpf, 72 hpf, and 120 hpf (n=10 for each stage and condition). (**G**) Stacked Histogram showing the number of neuromasts in the supraorbital region of DMSO control versus NIF treated specimens at 48 hpf, 72 hpf, and 120 hpf (n=10 for each stage and condition). Abbreviations are as follows:-gAD: anterodorsal lateral line ganglion; gAV: anteroventral lateral line ganglion; gP: posterior lateral line ganglion; hb: hindbrain. Data are shown as the mean ± SD (E, F, G) ns P > .05, *P < .05, ***P < 0.001 (Wilcoxon Test). Scale bars are 50 µM (A-C) and 20 µM (D).

In Figure 7 we compare confocal maximum projections of high magnification lateral views, showing the anterior and posterior lateral line ganglia and the afferent nerves of untreated control (Fig. 7A), DMSO-treated control (Fig. 7B), and NIF-treated experimental (Fig. 7C) 120 hpf *Tg(sox10:NTRmCherry;cntnap2a:EGFP)* double transgenic specimens. Compared with the two control conditions, which show no significant differences from one another, NIF-treated specimens lack most of the posterior lateral line ganglion (gP) cells, and have a significant reduction in the anterior lateral line ganglia. Moreover, the gAD and gAV components of the anterior lateral line ganglia can no longer be distinguished in experimental specimens (gAD/gAV, Fig. 7C). While the controls show distinct projections from the gAD and gAV into the hindbrain, NIF-treated specimens lack the gAV projection and have thinner, defasciculated nerves (compare Figs 7A, B, C, arrowheads). Fig. 7D shows an example of the defasciculation typical in neuromast-innervating nerves of neural crest-deficient specimens.

We next quantified aspects of these phenotypes. Fig. 7E is a Boxplot comparing counts of anterior lateral line ganglia neurons in DMSO-control specimens (blue) with NIF-treated specimens (red), revealing a significant reduction in number of neurons when neural crest cells are missing. Fig. 7F quantifies the number of supraorbital nerve projections that innervate neuromasts in DMSO-treated control specimens (blue) versus NIF-treated specimens (red) at 48 hpf, 72 hpf, and 120 hpf. This analysis reveals that when neural crest cells are lacking there is a modest increase in the number of nerve projections at 48 hpf and 72 hpf, followed by a significant increase at 120 hpf. The increase in projections correlates with the increase in neuromasts, as shown in Fig. 7G, which quantifies the number of neuromasts in the supraorbital region of DMSO-treated control specimens (blue) versus NIF-treated specimens (red) at 48 hpf, 72 hpf, and 120 hpf. When neural crest cells are missing there is an increase in the number of neuromasts in the supraorbital region, which becomes significant at 120 hpf.

Finally, to gain a more dynamic understanding of how inappropriately localized anterior lateral line neuromasts and disrupted nerves develop in the absence of neural crest cells, we again turned to SPIM time-lapse analysis. Figure 8 shows a series of still images taken from a time-lapse analysis of a NIF-treated *Tg(cldnB:GFP;sox10:NTRmCherry)* double transgeni*c s*pecimen, imaged between 33 and 55 hpf (Fig. 8A-F; and Supplemental Videos 5, 6). By 33 hpf (Fig. 8A) the supraorbital (SOp) and infraorbital (IOp) primordia had split anterior to the otic vesicle (ov). However, unlike unmanipulated control specimens with a full complement of neural crest cells (Figs 1, 2), in this specimen a broad region of Cldn:GFP labeled tissue (asterisk) remained between the supraorbital and infraorbital branches.

**Figure 8:**
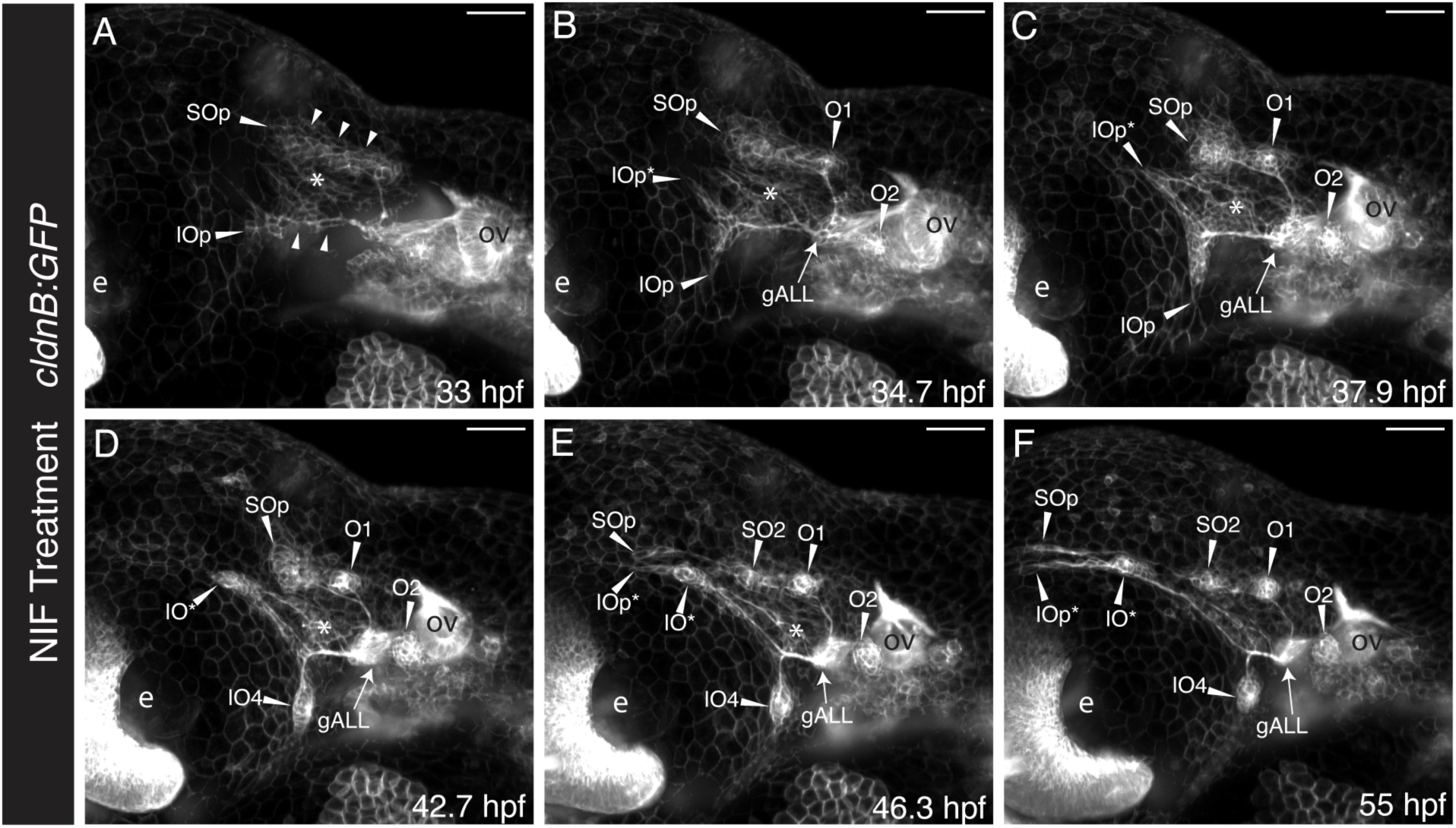
Time-lapse analysis of anterior lateral line development in the absence of neural crest cells. (**A-F**) SPIM time-lapse images of the cranial region of a *Tg(sox10:NTRmCherry;cldnB:GFP)* double transgenic specimen, treated with Nifurpirinol (NIF), in lateral view with anterior to the left. Single channel maximum projections, imaged using the 10x objective, showing only the *cldnB:GFP* label are shown at (A) 33 hpf, (B) 34.7 hpf, (C) 37.9 hpf, (D) 42.7 hpf, (E) 46.3 hpf, and (F) 55 hpf. (A) Arrowheads indicate primordia of the supraorbital (SOp) and infraorbital (IOp) lines splitting anterior to the otic vesicle at 33 hpf, as they begin to migrate. (B) shows the infraorbital primordium splitting into two primordia at 34.7 hpf, a ventrally projecting branch (IOp) and an ectopic anterodorsally projecting branch (IOp*); otic neuromasts O1 and O2 have formed. (C) shows that by 37.9 hpf the ectopic dorsal infraorbital primordium (IOp*) has migrated parallel to the supraorbital primordium (SOp), while the ventral infraorbital primordium (IOp) has migrated ventrally to produce infraorbital neuromast IO4. (E) by 46.3 hpf SOp has dropped off the first supraorbital neuromast (SO2) and continued to extend, but without producing an SO1 neuromast. The anterodorsal IOp* primordium has dropped off an ectopic neuromast IO* and continued to migrate anteriorly, parallel to SO1p. (F) shows that ectopic neuromast IOp* is innervated by a nerve from the anterior lateral line ganglion gALL. The ventral IOp* stalls after producing IO4 and does not continue to migrate below the eye. Abbreviations are as follows: e: lens of the eye, ov: otic vesicle. Scale bars are 50 µm.

By 34.7 hpf (Fig. 8B) the otic neuromasts O1 and O2 had condensed, while the supraorbital primordium SOp had continued to migrate dorsoanteriorly over the eye (e). Distinct from specimens with intact neural crest cells (Figs 1, 2 and Supplemental Movies 1-4), in this experimental specimen (Fig. 8B) the infraorbital primordium split into a dorsal branch (IOp*) as well as a ventral branch (IOp), as it approached the eye. By 37.9 hpf (Fig. 8C) the ectopic dorsal component of the infraorbital primordium (IOp*) had begun migrating parallel to the supraorbital primordium SOp, while the ventral component of the infraorbital primordium (IOp) behaved as in unmanipulated specimens, migrating ventrally. The ventral component of the primordium started to condense into the first infraorbital neuromast by 42.7 hpf (Fig. 8D, IO4), while the dorsal component (IOp*) continued to migrate parallel to the supraorbital primordium (Fig. 8D, SOp). By 46.3 hpf (Fig. 8E), migration of the ventral infraorbital primordium had completely stalled, with IO4 now in place. The supraorbital primordium deposited neuromast SO2, and its migration stalled at the apex of the eye (Fig. 8E, SOp). The dorsal branch of the infraorbital primordium (IOp*) similarly stalled parallel to SO1p at the apex of the eye, after having dropped off an ectopic neuromast IO* (Fig. 8E). By 55 hpf (Fig. 8F), the supraorbital and ectopic dorsal branch of the infraorbital primordia (SOp and IOp*) had ceased to migrate, after having deposited neuromasts SO2 and IO*, respectively. Axons can be seen extending from the anterior lateral line ganglia (gALL) to the otic neuromast O1, the supraorbital neuromast SO2, the infraorbital neuromast IO4, as well as ectopic neuromast IO* (Fig. 8F). The region of ectopic Cldn:GFP labeled tissue between the supraorbital and infraorbital branches (asterisks, Fig. 8A-E) gradually reduced in size over developmental time, consistent with the possibility that these cells might become incorporated into the ectopic dorsally projecting IO primordium (IOp*).

In summary, we found that when neural crest cells are absent, there are not only disruptions in neuromast migration, number, and organization, but also related disruptions in their innervation and reductions in the size of the anterior lateral line ganglion.

### 2.6 The most anterior neuromasts are encased in neural crest-derived bone in adult zebrafish

As zebrafish ontogeny proceeds, the lateral line system remains closely associated with neural crest cell-derivatives. Supraorbital canals, which will enclose the primary SO neuromasts, begin to develop in postembryonic specimens that are 10 mm standard length (around 4-6 weeks post fertilization; Parichy et al., 2009) (Webb & Shirey, 2003). By the time the developing fish are young adults of 22 mm in standard length, the process of canal enclosure of the neuromasts is close to complete (Webb & Shirey, 2003). Having observed early patterning defects in the most anterior components of the supraorbital system when neural crest cells were absent, we hypothesized that these same anterior components are the ones that ultimately lie in neural crest cell-derived bone.

The contributions of neural crest cells to the bones of the zebrafish cranium have previously been mapped out using the Cre-recombinase based two-transgene system described above (Kague et al., 2012). This analysis established that only the most anterior component of the frontal bone is neural crest derived, with the remainder being mesoderm-derived. The small nasal bone, which lies anterior to the frontal bone, is similarly neural crest derived (Kague et al., 2012). We exploited the same two-transgene Cre-based lineage tracing system to follow neural crest-derived cells into the bony elements of the cranium, to ask how these neural crest-derivatives correspond with supraorbital lateral line components.

We began our analysis by using *Tg(sox10Cre;dsRed/EGFP)* zebrafish to follow neural crest-derived cells into juvenile zebrafish. Fig. 9A shows a maximum projection confocal image of the left side of the cranium, in dorsal view, of a 6 mm standard length (SL) (around 3 weeks post fertilization) juvenile specimen. For this analysis, we labeled the supraorbital neuromasts using the vital mitochondrial dye TMRE (yellow), and found that the anteriorly localized SO1 neuromast was now embedded in neural crest-derived, *sox10Cre;EGFP*-positive (magenta), tissue. We then extended our analysis to young adult stages (22 mm SL). Fig. 9B shows a low-magnification maximum projection large format lightsheet image of the right side of the cranium of a *Tg(sox10Cre;dsRed/EGFP)* specimen. This specimen is co-labeled with a hair cell marker, anti-Otoferlin antibody (yellow) (Goodyear et al., 2010), to reveal SO neuromasts, which now include the newly formed anteriorly-located SO1’ neuromast (Webb & Shirey, 2003), as indicated. By this stage, the supraorbital canal has formed, and its large open pores are readily apparent, highlighted by edge effects that produce some non-specific fluorescent signal in these large specimens. To provide more context, Fig. 9C shows a schematic view of the entire cranial region of a 22 mm SL young adult specimen, again in dorsal view, indicating the supraorbital canals and the canal neuromasts within them (yellow), in relation to the neural crest-derived (magenta) and mesoderm-derived (blue) regions of the frontal bone. The small anterior nasal bones, which are also neural crest-derived, lie adjacent to the nostrils (nos). The region imaged in Fig. 9B-B’’ is indicated by a dashed box.

**Figure 9:**
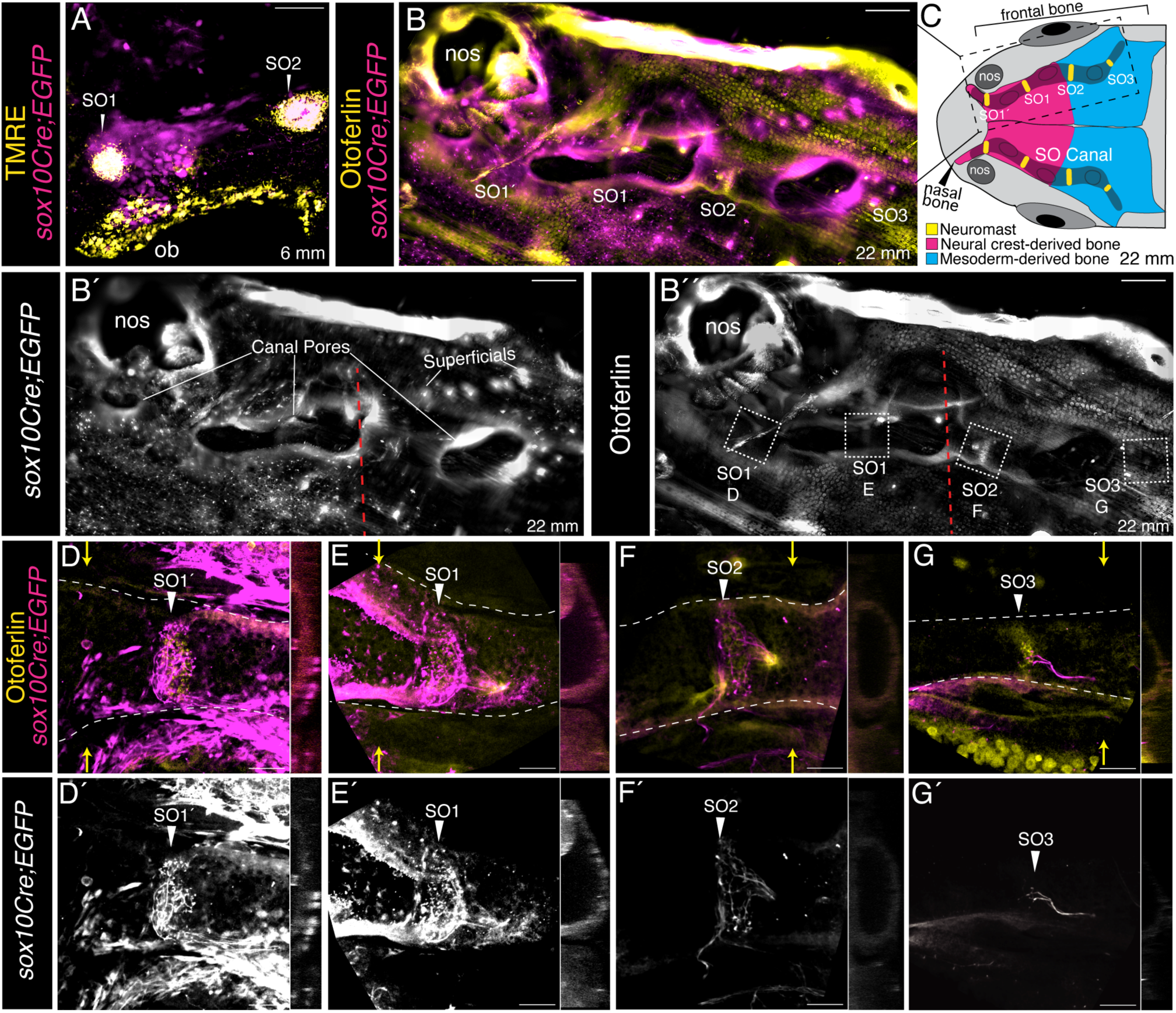
The most anterior supraorbital neuromasts are encased in neural crest-derived bone in adult zebrafish. (**A**) Confocal maximum projection (40x objective), in dorsal view, of the left hand side of the anterior cranial region of a 6 mm SL *Tg(sox10Cre;dsRed/EGFP*) specimen labeled with the vital dye TMRE, a mitochondrial marker labeling neuromasts. Canal neuromasts SO1 and SO2 are shown (TMRE, yellow) alongside neural crest-derived tissue (*sox10Cre;EGFP*, magenta). Because TMRE is highly fluorescent in the rhodamine channel (546 nm), the *sox10Cre:dsRed* signal present throughout the specimen is not visible with the imaging parameters selected. Ob = olfactory bulb. (**B**) Large format lightsheet (4x objective), maximum projection of the right hand side of the cranium (boxed area in C) of a 22 mm SL adult *Tg(sox10Cre;dsRed/EGFP*) specimen (of n=5). Neural crest cell-derived tissue is labeled with *sox10Cre;EGFP* (magenta), neuromasts are labeled with anti-Otoferlin antibody (yellow), in dorsal view with anterior to the left. (**B’**) EGFP channel showing neural crest cell-derived tissue. (**B’’**) Otoferlin channel showing neuromasts (in white dashed-line boxes). (**C**) Schematic of the supraorbital canals and canal neuromasts (yellow) in an adult zebrafish. The large frontal and small anterior nasal bones (next to the nostrils, nos) are indicated. Cell-type contributions to the bones are as indicated, neural crest in magenta, mesoderm in blue. (**D-G’**) Confocal maximum projection (40x objective), in dorsal view with anterior to the left, of each of the canal neuromasts shown in B’’ (boxed regions). The white dashed lines in D-G indicate the supraorbital canal; D’-G’ show the *sox10Cre;EGF*P channel alone. Orthogonal (yz) sections, taken adjacent to each neuromast (at level of yellow arrows), are provided to the right of each panel. Scale bars are 50 µm for panels A, D-G’’ and 200 µm for panels B-B’’.

The image in Fig. 9B is shown in single channels in Figs 9B’ and 9B’’. Confirming a previous report (Kague et al., 2012), Fig. 9B’ (the *sox10Cre;EGFP* channel) shows that the anterior part of the frontal bone is neural crest cell-derived; note EGFP-positive puncta across the entire surface of the region to the left of the red dashed line. Newly formed superficial neuromasts are also apparent at this late stage (Webb & Shirey, 2003). In common with the canal neuromasts (see below), the superficials include EGFP-positive puncta, reflecting innervation from neural crest cell-derived neurons. Fig. 9B’’(the Otoferlin channel) allows individual canal neuromasts to be identified. The region surrounding each of these neuromasts (white dashed line boxes) is re-imaged using confocal microscopy at high magnification in Fig. 9D-G’, with orthogonal (yz) views captured adjacent to each neuromast (plane indicated by yellow arrows), provided to the right of each image. Of note, at the level of each neuromast there are filamentous EGFP-positive structures, which are the tips of the neural crest cell-derived afferent nerves. Importantly, Figs 9D, D’ and E, E’ show that the bone surrounding supraorbital neuromasts SO1 and SO1’ is EGFP-positive, confirming its neural crest origin. By contrast, Figs. 9F, F’ and 9G, G’ show that the bone surrounding SO2 and SO3 lacks a neural crest contribution.

In summary, our Cre-based lineage tracing has shown that the most anterior components of the supraorbital lateral line system, the SO1 neuromasts and the late forming SO1’ neuromasts, lie in the neural crest-derived portion of the frontal bone and in the neural crest-derived nasal bone, respectively, while SO2 and SO3 are located in the mesoderm-derived portion of the frontal bone.

## 3. Discussion

Our study has augmented understanding of zebrafish anterior lateral line system development and its relationship with the cranial neural crest. Importantly, we have documented a close physical association between the developing placodally-derived anterior lateral line system and the underlying cranial neural crest cells. Further, we have tested the functional significance of this interaction. Using transgene mediated cytotoxicity to kill neural crest cells, we have demonstrated that the zebrafish cranial neural crest plays a key role in the development of the overlying anterior lateral line. Both the supraorbital and infraorbital lines fail to extend into the anterior, pre-orbital, portion of the cranium in the absence of neural crest. Using genetic lineage tracing, we have established that later in ontogeny it is these same anterior supraorbital neuromasts that become encased in neural crest-derived bone. Taken together, our experiments have revealed that the cranial neural crest and cranial placode derivatives function in concert over the extent of ontogeny to build the complex anterior lateral line system.

Our analysis of anterior lateral line development during the first five days of zebrafish larval life is broadly consistent with a previous comprehensive description of neuromast deposition (Iwasaki et al., 2020). Like Iwasaki and colleagues (2020), we have documented migrating anterior lateral line primordia, neuromast budding processes, and the emergence of intercalary neuromasts from interneuromast cells. However, our SPIM time-lapse analysis has further revealed that migration and budding in the supraorbital line are not entirely separate processes, with the budding of SO1 from SO2 beginning before SO2 has reached its final location (Fig. 1; Supplemental Videos 1-4). Moreover, our Single Plane Illumination Microscopy reveals that the infraorbital line initially forms as an elongated ridge of cells, with neuromasts subsequently emerging through condensation and proliferation of progenitor cells (Fig. 1; Supplemental Videos 3, 4). The existence of this ridge was not noted by Iwasaki et al. (2020), possibly reflecting the challenge of imaging cells that are located immediately below the developing orbit using standard confocal microscopy.

Migration of a primordium that deposits neuromasts in its wake has been described in great detail in the context of the zebrafish posterior lateral line, and is thus the more familiar process, but it is important to note that this might not be the general condition in jawed vertebrates. Rather, elongation and fragmentation of lateral line primordia occur in both crania and trunks of chondrichthyan species (Gillis et al., 2012; Johnson, 1917), as well as in the crania of many non-teleost osteichthyan species (Modrell et al., 2011; Northcutt et al., 1994; Stone, 1928; Winklbauer & Hausen, 1983). As noted by Iwasaki et al. (2020) evolution of different modes of neuromast deposition might reflect relatively modest changes in specific cellular properties. Ultimately, a full understanding of lateral line system evolution will require documentation of these varying deposition mechanisms across the vertebrates, including in a wider range of teleosts.

Our analysis of lateral line gangliogenesis and afferent nerve development is in agreement with the seminal study of Raible and Kruse (2000), as well as with the recent detailed transgenic analysis of Iwasaki and colleagues (2020). We focused in particular on the relationship of developing lateral line ganglia and nerves with the neural crest cells. Confirming a report from Kague et al. (2012) we documented that the neurons in the anterior and posterior lateral line ganglia derive from both placodes and neural crest. We extended their findings to confirm that both the dorsal (gAD) and the later developing ventral (gAV) components of the anterior lateral line ganglia include neurons of both origins. We also established that when the lateral line ganglia first develop from the placodes they are surrounded by a ‘shell’ of neural crest cells, with other neural crest cells—presumably developing glia—interacting closely with the developing anterior lateral line nerves.

When neural crest cells were removed by transgene-mediated cytotoxicity we found that anterior lateral line development was dramatically disrupted. In the absence of neural crest cells the migration of the supraorbital primordium ‘stalls’ above the apex of the eye, such that no anterior supraorbital neuromasts are deposited in their typical anterodorsal locations relative to the eye. Similarly, the infraorbital primordium fails to extend ventrally below the eye, leading to a similar absence of anterior neuromasts in their typical anteroventral locations relative to the eye. In addition, supernumerary neuromasts often develop in the absence of neural crest, producing an unusually condensed array of neuromasts in the posterior supraorbital region. Potential sources of these supernumerary neuromasts are discussed below.

As neural crest cells contribute directly to the lateral line ganglia, it is not surprising that we find alterations in both gangliogenesis and lateral line innervation when neural crest cells are missing. While not a primary focus of this study, we note that the posterior lateral line ganglion (gP) is severely reduced in size and highly disorganized in the absence of neural crest cells (as exemplified by Figure 7C). Despite this, we find that the posterior lateral line primordium migrates successfully to the terminus of the trunk, consistent with previous reports that innervation is not required for its migration (López-Schier & Hudspeth, 2004). While the anterior lateral line ganglion shows less dramatic deficits, our quantification reveals that the number of neurons in the ganglion is significantly reduced by 5 days post fertilization, with the anteroventral (gAV) component of the ganglion either absent or drastically reduced in size, and the projection from gAV into the hindbrain lacking altogether (Figure 7). Of note, it is this late developing gAV component of the ganglion that innervates the mandibular and opercular lines, which are completely missing when neural crest is absent, presumably reflecting the absence of the tissue in which they would normally form.

In addition to reduced gAD ganglion size when neural crest cells are absent, the anterior lateral line nerves are thin, defasciculated, and often make ectopic projections (Figures 6, 7). These ectopic projections frequently, but not always, connect with ectopic neuromasts in the supraorbital region. On occasion, the nerve that typically projects anteroventrally to the infraorbital line (nADio) instead—or in addition—projects more dorsally, raising the possibility of dorsal translocation of a single IO neuromast into the posterior supraorbital domain.

Consistent with this hypothesis, our SPIM time-lapse analysis (Figure 8; Supplemental Videos 5, 6) has captured an example of such aberrant dorsal migration. Thus, at least in some instances, ectopic neuromasts in the supraorbital region might be misplaced infraorbital neuromasts that are following supraorbital cues and migrating above the eye. Overall, the deficits we have found in lateral line gangliogenesis and innervation are consistent with previous descriptions of roles for neural crest in ganglion condensation and establishment of neural connections (reviewed by Steventon et al., 2014), including a requirement for chondrogenic—but not glial— neural crest in the initial assembly of zebrafish epibranchial ganglia (Culbertson et al., 2011).

Our results raise questions about the source of the other ectopic neuromasts in the posterior supraorbital region of neural crest-deficient zebrafish larvae. One possibility is that the ectopic neuromasts derive from cells which would normally form the mandibular or opercular neuromasts. Another possibility, by analogy with posterior lateral line phenotypes in the absence of neural crest-derived glia, is that these additional cranial neuromasts represent prematurely or ectopically developing intercalary neuromasts, or both. We have observed neural crest cells in close proximity to the projecting supraorbital nerves (Figures 2, 3), and these ‘wrapping’ cells are presumably myelinating Schwann cells (Lyons et al., 2005). In the absence of the Schwann cells, nearby interneuromast cells might proliferate precociously and condense to form new neuromasts, much as occurs in the trunk. However, it should be noted that a previous study of *ngn1* and *sox10/colourless* mutants, both of which lack glial cells, did not reveal ectopic cranial neuromasts at 6 days post fertilization (López-Schier & Hudspeth, 2005). Nevertheless, a more detailed future exploration of a potential role for Schwann cells in suppressing intercalary neuromast development in the anterior lateral line system is warranted. If Schwann cells prove to play non-equivalent roles in the cranium and trunk, with respect to suppressing precocious neuromast formation, this does not rule out a similar role for other cranial neural crest-derived cells.

We have found that both the neuromasts and the lateral line nerves show disrupted localization in the absence of neural crest. While the anterior lateral line neuromasts and their nerves develop together and in concert in specimens with intact neural crest, we have found that when neural crest cells are lacking the lateral line nerves project from non-uniform points on the ganglia, and then tend to wander, not always innervating a target neuromast. These observations suggest that neural crest cells influence nerve outgrowth from the ganglia. It remains to be resolved whether the relevant neural crest cells lie in the outlying tissue, wrap around the ganglia, contribute neurons to the ganglia, or are in some combination of these locations. Subsequent axon guidance may also be influenced by neural crest-derived cells, particularly glia. Consistent with such a model, it has been shown that glial-derived neurotrophic factor (GDNF) is required for proper extension of the posterior lateral line nerve, although in that instance it is the posterior lateral line primordium itself that serves as the GDNF source (Thomas et al., 2015).

The precise nature of the interaction between neural crest and placode-derived lateral line cells deserves further attention. While the neural crest is a highly migratory cell type, our time-lapse analyses reveal that neural crest cells neither tow nor chase (Thevenau et al., 2013) the anterior lateral line primordia. Rather the neural crest cells are already in place, although still proliferating and far from static (Supplemental Videos 2 and 4), at the stages when the supraorbital and infraorbital primordia are migrating over them. The neural crest cells might, however, be providing a physical substrate on which the anterior lateral line primordia migrate and bud, and our confocal analysis has revealed close apposition of the two cell types, compatible with such a model. Alternatively, the neural crest cells might serve as the source of one or more localized signals. Available data (reviewed by Piotrowski & Baker, 2014) suggest that anterior lateral line development does not require chemokine signaling—in distinct contrast to the situation in the posterior lateral line—but one or more different short-range signals could potentially be at work.

Iwasaki et al. (2020) uncovered a role for local tissue-tissue interactions in the specific instance of hyomandibular neuromast progenitors; these respond to R-spondin2 emanating from mesenchyme of the second pharyngeal arch, which in turn activates Wnt/beta-catenin signaling to activate progenitor proliferation. The lack of a requirement for R-spondin2 in the development of other cranial neuromasts is consistent with a model in which multiple different localized molecular signals, or cues, serve to pattern the varied components of the anterior lateral line system. We hypothesize that one such cue (or set of cues) is followed by the supraorbital primordium as it initially migrates dorsoanteriorly to the position where SO2 is deposited. The relatively normal initial migration of the supraorbital primordium in the absence of neural crest cells suggests that this particular cue must emanate from non-neural crest derived tissue. Moreover, we observed that when neural crest cells are missing, a dorsally mis-placed infraorbital primordium can migrate adjacent to the supraorbital primordium, presumably following the supraorbital cue(s) (Figure 8; Supplemental Movies 5, 6). A candidate for this supraorbital cue is Fgf signaling. Simultaneous knockdown of Fgf3 and Fgf10a disrupts supraorbital neuromast deposition, with the sources of the Fgfs identified as the endoderm and the anterior lateral line itself, respectively (McCarroll & Nechiporuk, 2013).

Our study has focused on the anterior lateral line system, which was already understood to use distinct developmental mechanisms relative to those of the better studied posterior lateral line system (Piotrowski & Baker, 2014). However, our data have uncovered another important difference between the head and trunk lines: we have demonstrated that in the absence of neural crest cells the anterior lateral lines fail to reach their destinations, yet the posterior lateral line completes its full migration to the caudal terminus of the trunk (Figure 5).

Intriguingly, the posterior lateral line system, despite its destination being the trunk, is nevertheless derived from a cranial placode, and could be viewed as a “head” structure (Northcutt & Gans, 1983). Nevertheless, this post-otic placode derivative responds very differently to adjacent neural crest cells than does the pre-otic anterior line placode. It remains unclear to what extent these differences are driven by intrinsic differences between the placodes, for example differential Hox gene expression status, versus extrinsic differences.

Classical embryology experiments performed by Stone (1928) in the axolotl suggested that extrinsic differences play a major role. Stone swapped the anterior and posterior lateral line placodes, and found that these swapped placodes responded much as the correct ones to their new environments. The powerful molecular genetics and imaging opportunities now available in the zebrafish provide opportunities to continue to explore the molecular underpinnings of both intrinsic and extrinsic aspects of lateral line patterning, including the bases of the differences between the anterior and posterior systems.

The placode-derived lateral line system and the neural crest cells are two vertebrate specific cell types that are key to vertebrate evolution (Gans & Northcutt, 1983; Hall, 1998; Northcutt & Gans, 1983). The relationship between these cell types begins at the earliest developmental stages with their shared embryonic origins from nearby or overlapping non-neuroepithelial ectoderm (reviewed by Koontz et al., 2023; Rocha, Beiriger, et al., 2020; Rocha, Singh, et al., 2020; Steventon et al., 2014). The relationship continues with multiple functional interactions, now including those we describe in this study in which the neural crest cells influence anterior lateral line patterning. This relationship then culminates with elements of the lateral line system becoming embedded in the dermal bone that derives from this same neural crest population (this study, Webb & Shirey, 2003). The contributions of neural crest to different components of the cranial dermal skeleton have changed through the course of vertebrate evolution (Rocha, Beiriger et al., 2020), as have the morphologies of lateral lines (Northcutt, 1989). New comparative data will be needed to elucidate how the relationship between placode-derived lateral lines and cranial neural crest has been modified over evolutionary time scales to produce the numerous different lateral lines patterns that we find in different vertebrate groups (Northcutt, 1989).

As discussed in the introduction, Parrington (1949) hypothesized that the precursors of dermal ossifications—which we now know include neural crest cells—might influence the ‘courses’ of the lateral lines. Parrington also noted that the test of his hypothesis must “rest eventually on experimental evidence”. Three-quarters of a century later, our study now provides that evidence, validating Parrington’s insightful model and confirming that there is an intimate and ongoing functional relationship between the cranial neural crest cells (which include precursors of dermal bones) and placode-derived anterior lateral lines.

## 4. Materials and Methods

### 4.1. Animal Husbandry

Zebrafish (*Danio rerio*) were maintained in accord with IACUC-approved protocols at the University of Chicago, the Stowers Institute, and the Marine Biological Laboratory. Embryos were maintained in E3 solution (5mM NaCl, 0.17mM KCl, 0.33mM Ca_2_Cl_2_, 0.33mM MgSO_4_), which was supplemented with 0.3% 1-phenyl-2-thiourea (PTU; Sigma) to block pigment formation for analyses after 24 hpf, and staged according to standard guidelines (Kimmel et al., 1995; Parichy et al., 2009). Embryos were obtained from crosses of adult fish stocks of wild types (*AB line) and/or transgenics.

The following transgenic zebrafish lines were used in this study: *Tg(cldnB:GFP)^zf106^*, a tight junction marker that labels the membranes of cells in the lateral line system, other placodes, and epithelia (Haas & Gilmour, 2006); *Tg(−8.0cldnB:H2AmCherry)^psi4Tg^*, referred to in the text as *Tg*(*cldnB:H2AmCherry)*, is a nuclear localized red fluorescent version of the same marker (Peloggia et al., 2021); *Tg(dRA:GFP)*, a fortuitous insertion line that labels the lateral line primordia (provided by Parker and Krumlauf, who generated the line using methodology described in (Parker et al., 2014)); *Tg(−7.2sox10:mRFP)^vu234^*, referred to in the text as *Tg(sox10:mRFP)*, labels the membranes of neural crest cells as well as part of the otic vesicle (Kucenas et al., 2008)*; Tg(sox10:gal4;UAS:NTR-mCherry)*, which combines the *Tg(−7.2sox10:Gal4-VP16)^psi5^* (Rosenberg et al., 2014) and *Tg(UAS-Eib:NfsB-mCherry)^jh17^* (Parsons et al., 2009) transgenes, this line is referred to in the text as *Tg(sox10:NTRmCherry)*, and expresses Nitroreductase enzyme in neural crest cells and the otic vesicle; *Tg(Hgn39d:EGFP)* is an enhancer trap insertion (Nagayoshi et al., 2008) into the *contactin associated protein 2a* gene, referred to in the text as *Tg(cntnap2a:EGFP)*, and exclusively labels lateral line afferent nerves (Faucherre et al., 2009; Pujol-Martí et al., 2012); *Tg(−28.5Sox10:cre;ef1a:loxP-dsRed-loxP-egfp*), referred to in the text as *Tg(sox10Cre;dsRed/EGFP*), uses Cre/Lox recombination to permanently label neural crest cells with EGFP, while other cells express dsRed (Kague et al., 2012).

### 4.2. Single Plane Illumination Microscopy (SPIM)

Transgenic embryos were mounted for SPIM in Fluorostore Fractional FEP Tubing (F018153-5) using a modified multilayer technique (Kaufmann et al., 2012). Embryos were immobilized using 0.3% agarose (Invitrogen UltraPure Low Melt Agarose Cat #16500) dissolved in E3 medium and 0.2 mg/ml tricaine, and the FEP tubing capped with a 1.2% agarose plug. Embryos were incubated at 28.5 °C during data collection. Images were captured with either a Zeiss Lightsheet Z.1 (with 20x objective) or a Zeiss Lightsheet 7 (with 10x objective) single-plane illumination microscope, each equipped with tandem PCO.edge sCMOS cameras (PCO.Imaging, Kelheim, Germany). Volumes were acquired every 10 minutes. Zeiss Zen software was used to acquire images, with post processing in FIJI (Schindelin et al., 2012) and Imaris (Oxford Instruments).

### 4.2. Confocal image acquisition

For assays in fixed specimens, embryos were fixed in 4% paraformaldehyde (PFA; Sigma) at 4 °C overnight. Following overnight fixation, embryos were washed in 1X Phosphate Buffered Saline (PBS) five times for 5 min each. For long-term storage of embryos, embryos were washed in 30%, 60% and 100% methanol (diluted in 1X PBS) and stored in 100% methanol at −20 °C. If stored in 100% methanol, embryos were progressively washed in 60%, 30% methanol as well as 1X PBS + 0.1%Tween-20 (Sigma) before mounting or staining. Static confocal images were collected on upright Zeiss LSM710 or LSM900 confocal microscopes with Plan-Apochromat 10x/0.45, 20x/0.8W, or 40x/1.W objectives. Time-lapse confocal microscopy of living specimens was performed on an inverted Zeiss 780 microscope using a 40x/1.1W Corr M27 objective in a climate-controlled chamber set at 28 °C. Embryos were anesthetized as described for SPIM and mounted in 0.8% low melt agarose in glass-bottomed dishes (MatTek, USA). Zeiss Zen software was used to acquire images, with post processing in FIJI (Schindelin et al., 2012).

### 4.3. Nitroductase mediated neural crest ablations

Double transgenic zebrafish specimens were acquired by crossing individuals of two different transgenic lines to generate *Tg(sox10:NTRmCherry;cldnB:GFP)* or *Tg(sox10:NTRmCherry;cntnap2a:EGFP*) embryos. The *sox10:NTRmCherry* transgene is expressed shortly after 9 hours post fertilization, so drug treatments were begun at this stage. To achieve drug-mediated ablation of Nitroreductase-expressing neural crest cells, 9 hpf embryos were transferred in their chorions into 1.25 μM Nifurpirinol (Sigma-Aldrich catalog # 32439) dissolved in E3 medium plus 1:1000 dimethyl sulfoxide (DMSO) and incubated in the dark. Controls were incubated in E3 medium plus DMSO carrier alone. Double transgenic embryos were sorted and dechorionated at 24 hpf, and single transgenic sibling specimens lacking *Tg(sox10:NTRmCherry)* also kept as controls. The Nifurpirinol solution was replaced with fresh solution at the 48 hpf stage for specimens to be raised to later stages.

### 4.4. Immunolabeling

Immunolabeling was performed as previously described (Prince et al., 1998) on specimens fixed in 4% PFA. The following primary antibodies were used: anti-Sox2 1:200 (anti-rabbit; GTX124477), anti-Sox10 1:200 (anti-rabbit; GTX128374), anti-HuC/D 1:200 (anti-mouse; Molecular Probes 16A11), anti-GFP 1:200 (anti-rabbit; Sigma A-6455), and anti-Otoferlin, 1:50 (anti-mouse; Developmental Studies Hybridoma Bank HCS-1). Secondary antibodies used were Alexa 488, 1:500 (Invitrogen anti-mouse A-11001; anti-rabbit A-11008), Alexa 546 1:500, (Invitrogen anti-rabbit A-11035), and Alexa 633, 1:300 (Invitrogen anti-mouse A-21052; anti-rabbit A-21070). The far red Alexa 633 secondaries were coupled with HuC/D, Otoferlin and Sox2 antibodies.

For adult zebrafish staining, a CUBIC clearing and staining protocol was adapted from Pende et al. (2020). Specifically, we depigmented adult specimens in acetone overnight at -20 °C before lightly bleaching for 5-19 minutes in a 3% solution of hydrogen peroxide in 1% KOH. Specimens were then incubated overnight in Low Urea CUBIC I solution at 37°C for initial clearing. For immunolabeling, cleared specimens were blocked using goat serum at 25°C for 3-4 hours before primary and secondary incubation. Specimens were incubated in primary antibody at 37°C for 2-3 days and subsequently in secondary antibody at 37 °C for 2 days, with *sox10Cre;EGFP* signal amplified using anti-GFP antibody. Specimens were placed in CUBIC R + (N) RI matching solution (Kubota et al., 2017) for at least 30 minutes prior to imaging. For TMRE live-dye labeling, larval zebrafish were incubated for 30 minutes in 5nM Tetramethylrhodamine, Ethyl Ester, Perchlorate (TMRE; Invitrogen) in E3 medium (Esterberg et al., 2013; Mandal et al., 2021).

### 4.5 Large format Lightsheet microscopy

Adult specimens were imaged on a LaVision Large Format Lightsheet, using a 4x objective, and 2.0 μM excitation sheet in CUBIC R+ (N) imaging medium (Kubota et al., 2017).

## Acknowledgements

We thank Tatjana Piotrowski for providing the *Tg(sox10:gal4;UAS:NTR-mCherry)* and *Tg(cldnB:GFP)* transgenic lines and for helpful discussions at the inception of the project. We thank Katie Kindt for providing the *Tg(Hgn39d:EGFP)* enhancer trap line, Lindsey Barske for providing the *Tg(−28.5Sox10:cre;ef1a:loxP-dsRed-loxP-egfp*) line, and Hugo Parker and Robb Krumlauf for providing the *Tg(dRA:GFP)* insertion line. We thank Ryan Anderson for invaluable advice on use of nitroreductase lines, and for sharing Nifurpirinol reagent with us. We are grateful to Karen Echeverri for generously making her space and equipment available to us during our visits to the Marine Biological Laboratory, and to Andrew Gillis and Cliff Ragsdale for providing helpful comments on the manuscript. We thank Adam Kuuspalu, Elaine Kushkowski, Michael Wen, Chris Bjornsson, Carsten Wolf, and Christine Labno for advice and assistance with imaging and image processing. Finally, we thank Adam Kuuspalu, Vanity Spruill, and Troy McInerney for expert zebrafish care.

## Supplemental Movies

**Supplemental Movie 1:** Confocal microscopy time-lapse movie of the cranial region of the *Tg(cldnB:GFP;cldnB:H2A-mCherry;dRA:GFP)* triple transgenic specimen shown in Figure 1A-E, in lateral view with anterior to the left. A single channel for the lateral line markers *cldnB:GFP* and *dRA:GFP* only is shown in white. Maximum projections from 25 hpf to 42 hpf.

**Supplemental Movie 2:** Same as Movie 1, with *cldnB:GFP* and *dRA:GFP* signals (yellow) and *cldnB:H2A-mCherry* nuclear label of lateral line and ectodermal epithelium (magenta) merged.

**Supplemental Movie 3:** SPIM time-lapse movie of the cranial region of the *Tg(sox10:NTRmCherry;cldnB:GFP)* double transgenic specimen shown in Figure 1F-I, in lateral view with anterior to the left. Only the *cldnB:GFP* lateral line marker (white) is shown. Maximum projections from 33 hpf to 52 hpf.

**Supplemental Movie 4:** Same as Movie 3, with the *cldnB:GFP* lateral line marker (yellow) and the neural crest cell marker (*sox10:NTRmCh*; magenta) merged.

**Supplemental Movie 5:** (**A-F**) SPIM time-lapse movie of the cranial region of the *Tg(sox10:NTRmCherry;cldnB:GFP)* double transgenic specimen treated with 1.25 µm Nifurpirinol (NIF) shown in Figure 8, in lateral view with anterior to the left. Only the *cldnB:GFP* lateral line marker (white) is shown. Maximum projections from 33 hpf to 55 hpf.

**Supplemental Movie 6:** Same as Movie 5, with the *cldnB:GFP* lateral line marker (yellow) and the neural crest cell marker (*sox10:NTRmCh*; magenta) merged. Note punctate dying neural crest cells (magenta).

